# Nuclear basket proteins regulate the distribution and mobility of nuclear pore complexes in budding yeast

**DOI:** 10.1101/2023.09.28.558499

**Authors:** Janka Zsok, Francois Simon, Göksu Bayrak, Luljeta Isaki, Nina Kerff, Yoana Kicheva, Amy Wolstenholme, Lucien E. Weiss, Elisa Dultz

## Abstract

Nuclear pore complexes (NPCs) mediate all traffic between the nucleus and the cytoplasm and are among the most stable protein assemblies in cells. Budding yeast cells carry two variants of NPCs which differ in the presence or absence of the nuclear basket proteins Mlp1, Mlp2 and Pml39. The binding of these basket proteins occurs very late in NPC assembly and Mlp-positive NPCs are excluded from the region of the nuclear envelope that borders the nucleolus. Here, we use recombination-induced tag exchange (RITE) to investigate the stability of all the NPC subcomplexes within individual NPCs. We show that the nuclear basket proteins Mlp1, Mlp2 and Pml39 remain stably associated with NPCs through multiple cell-division cycles, and that Mlp1/2 are responsible for the exclusion of NPCs from the nucleolar territory. In addition, we demonstrate that binding of the FG-nucleoporins Nup1 and Nup2 depletes also Mlp-negative NPCs from this region by an independent pathway. We develop a method for single NPC tracking in budding yeast and observe that NPCs exhibit increased mobility in the absence of nuclear basket components. Our data suggest that the distribution of NPCs on the nucleus is governed by multiple interaction of nuclear basket proteins with the nuclear interior.

**Significance Statement:**

- A subset of yeast nuclear pore complexes have a basket structure at their nuclear face. The stoichiometry, architecture and molecular functions of the basket are not clear.
- The authors use a tag exchange method to show stable binding of nuclear basket proteins through multiple generations. They also find that multiple nuclear basket proteins disfavour localization of NPCs proximal to the nucleolus.
- These findings demonstrate that having a nuclear basket is a final and stable state for yeast NPCs. The methods developed can be applied to gain more insight into the properties of NPCs and other stable complexes as they age.

## Introduction

Nuclear pore complexes (NPCs) are large macromolecular assemblies that form channels in the nuclear envelope (NE) and mediate all transport between the nucleus and cytoplasm in eukaryotic cells (reviewed in Dultz et al., 2022; Petrovic et al., 2022). The NPC is composed of multiple copies of approximately 30 different proteins, termed nucleoporins (Nups). Its core consists of a stack of three rings with eight-fold rotational symmetry along the nucleocytoplasmic axis and two-fold symmetry along the NE axis. The outer rings, on both the nucleoplasmic and cytoplasmic side of the NPC, are decorated with asymmetrical appendages known as the nuclear basket and the cytoplasmic export platform, respectively (Figure 1A).

**Figure 1.**
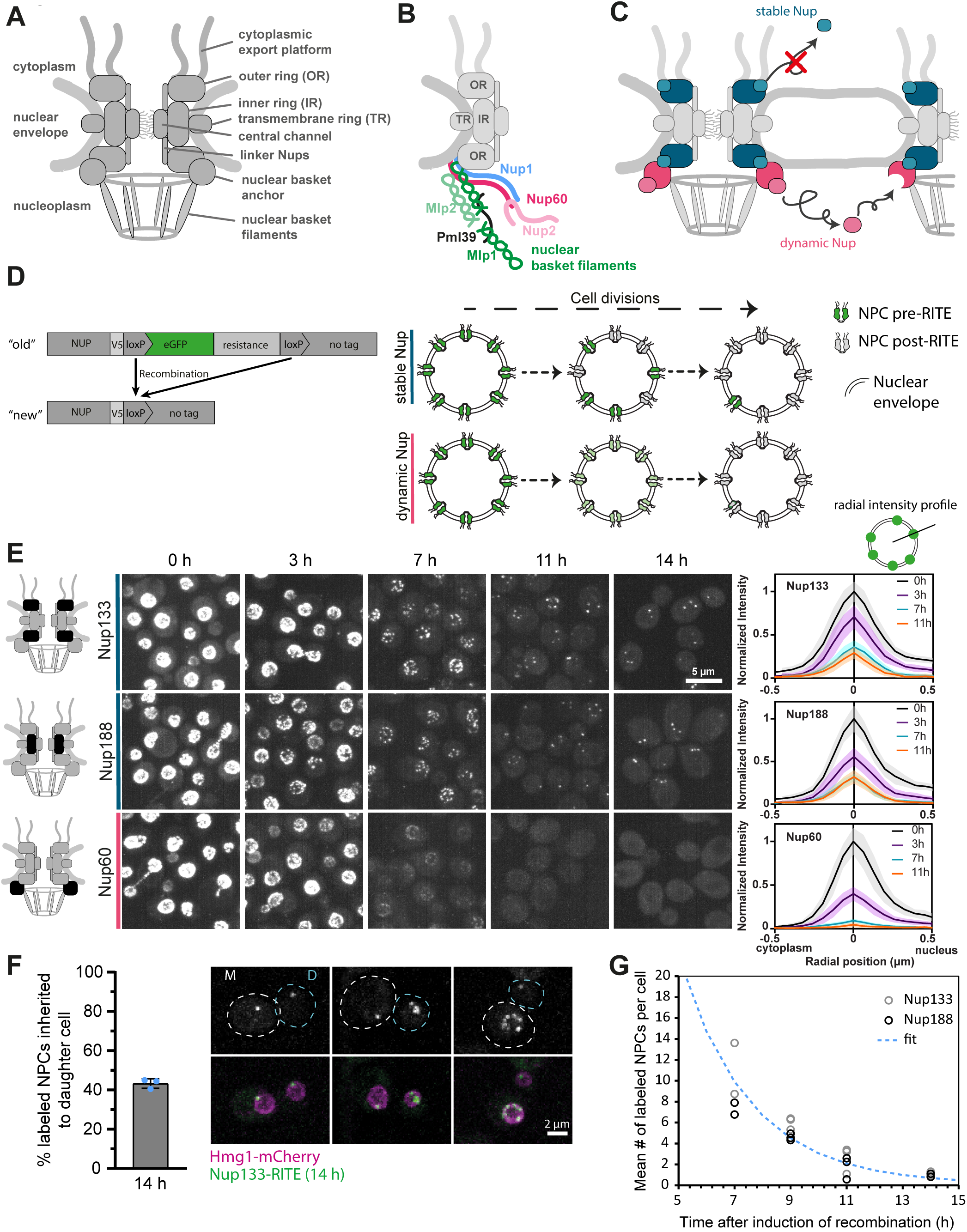
Single NPCs can be visualized using doRITE. (A) Overview of the subcomplexes of the NPC. (B) Schematic of the organization of the nuclear basket. OR (outer ring), IR (inner ring), TR (transmembrane ring). Nup60 anchors Nup2 and Mlp1 to the NPC. Mlp1 in turn is required for binding of Mlp2 and Pml39. Additional copies of Mlp1 are anchored by Pml39. (C) Behavior of a stable and a dynamic Nup. Stable Nups with a long residence half-life do not exchange with soluble subunits (blue). Dynamic Nups have a short residence half-life and readily exchange with a soluble pool and between NPCs (magenta). (D) Left: Schematic of the GFP-to-dark RITE cassette. NUP of interest is tagged with a V5 tag followed by a loxP site and eGFP (enhanced green fluorescent protein). Recombination leads to excision of the eGFP, so that only protein produced prior to recombination can be visualized as eGFP-tagged. Right: Schematic of the doRITE assay showing dilution of labelled Nups through cell divisions for a stable or dynamic Nup in yeast. (E) Dilution of RITE(GFP-to-dark)-labelled Nups through multiple cell divisions. Cells expressing the indicated Nups endogenously tagged with the RITE cassette were imaged at different timepoints after induction of RITE. Maximum projections are shown. All images are adjusted to the same brightness and contrast values. Colour bars represent membership to dynamic group in Figure 2A (blue: stable, magenta: dynamic). Schematics on left show location of Nup within the NPC. Graphs on right show radial intensity profile through fluorescent foci on the NE (compare schematic drawing on top) after indicated times of doRITE (Solid lines show mean of 20 cells, normalized for 0 h-timepoint, shaded area represents the standard deviation). (F) Inheritance frequency of RITE(GFP-to-dark)-labelled NPCs from mother (M) to daughter (D) cells after 14 h of doRITE. Number of NPCs was counted in 100 M-D pairs with 1-14 pores per pair. Bar graph shows mean of three biological replicates (mean of individual replicates are shown as blue dots), error bar represents the standard deviation. Three examples of M-D pairs are shown as maximum projections. Hmg1-mCherry marks the NE. (G) Mean number of labelled NPCs per cell determined from 100 cells per biological replicate at different time points after RITE(GFP-to-dark) induction for Nup133 or Nup188 (circles). The dashed line represents a fit of all shown datapoints with an exponential decay curve applying the experimentally determined doubling time of 1.75 h.

The nuclear basket is composed of three highly dynamic Nups carrying FG or FXF(G) domains (Nup1, Nup60 and Nup2) and three structural Nups (Pml39, Mlp1 and Mlp2), here collectively referred to as the basket filaments. The paralogues Mlp1 and Mlp2 are large proteins interspersed with coiled-coil regions that extend into the nucleoplasm, forming the main scaffold of the basket (Strambio-de-Castillia et al., 1999) (Figure 1B). They, along with Nup2, are recruited to the NPC by Nup60 via short linear interacting motifs (Dilworth et al., 2001; Cibulka et al., 2022; Stankunas & Köhler, 2024). Pml39 binds to Mlp1 and to a lesser extent to Mlp2 and promotes the binding of additional copies of Mlp1 to the NPC (Gunkel et al., 2023).

While Nup1, Nup60 and Nup2 are found on all NPCs, the nuclear basket filaments are present only on a subset (Galy et al., 2004; Palancade et al., 2005; Bensidoun et al., 2022). This may be a consequence of their late assembly: they bind to the NPC on average ∼40 min after all other Nups have assembled (Onischenko et al., 2020). The mechanism of this late assembly is unclear, but mRNA metabolism may be required for basket assembly, since inhibiting transcription and mutating mRNA maturation factors involved in 3’ end processing results in defects in basket filament assembly (Bensidoun et al., 2022). Supporting this idea, Mlp-positive NPCs are normally not found in the region of the NE adjacent to the nucleolus (Galy et al., 2004), which is devoid of mRNA. Mutants that accumulate mRNA in the nucleolus no longer exclude Mlp-positive NPCs from the nucleolar region (Bensidoun et al., 2022). There is thus an intricate relationship between the nuclear basket filaments and RNA maturation and export. Indeed, in wild type cells, mRNA export and processing factors are specifically enriched at NPCs with nuclear basket filaments (Bensidoun et al., 2022), and the deletion of *PML39* or *MLP1* results in the leakage of unspliced RNAs into the cytoplasm, suggesting that the nuclear basket plays a role in mRNA quality control (Galy et al., 2004; Palancade et al., 2005).

In addition to being involved in mRNA quality control, the nuclear basket filaments also contribute to genome protection by supporting DNA repair pathways and preventing R loops (Palancade et al., 2007; García-Benítez et al., 2017). Intriguingly, a recent study showed that telomeric gene silencing is specifically promoted by NPCs without nuclear basket filaments (Kumar et al., 2023). This is not the only example of NPC specialization, and there is increasing evidence that different NPC compositional variants carry out specific roles in the cell (reviewed in Dultz et al., 2022). For example, components of the mRNA export machinery are lacking in NPCs of daughter cells, which delays their transition into S phase (Gomar-Alba et al., 2022). Also, replicatively aged yeast cells accumulate NPCs that lack components of the cytoplasmic export platform and the nuclear basket, thereby contributing to ageing (Meinema et al., 2022). The study of such variant NPCs is severely limited by average-based bulk methods, since they fail to capture the compositional heterogeneity in different complexes. Thus, methods that can distinguish between NPC subtypes and provide spatial and temporal resolution of single NPCs are required.

In this work, we develop recombination-induced tag exchange (RITE) into a system that allows us to track individual NPCs and follow them through multiple cell cycles. We use this to systematically analyse the binding stability of Nups from all NPC subcomplexes in yeast. We show that Mlp1, Mlp2 and Pml39 are stably associated with NPCs through multiple cell cycles. Furthermore, we identify two pathways that reduce the density of NPCs in the nucleolar territory: First, binding of Mlp1/2 to mature NPCs prevents their localization in the nucleolar territory. Second, Nup2 leads to an increase of the residence time of all NPCs in the non-nucleolar territory, possibly by mediating interactions with chromatin.

## Results

### doRITE allows the visualization of individual NPCs in living budding yeast cells

Live-cell imaging of individual fluorescently tagged NPCs in budding yeast cells is severely limited by the high NPC density in the NE (>10 NPC/µm^2^) (Winey et al., 1997). To enable the visualization of individual NPCs, we reduced the density of fluorescently labelled NPCs on the nucleus by recombination-induced tag exchange (RITE) (Verzijlbergen et al., 2009). In RITE, a protein tag is genomically exchanged through Cre-induced recombination, so proteins produced before or after recombination can be distinguished by their different tags (Figure 1D). Because core Nups remain associated with individual NPCs over hours or even days (Hakhverdyan et al., 2021; Rabut et al., 2004), RITE can be used to distinguish between old and new NPCs (Colombi et al., 2013; Toyama et al., 2019; Onischenko et al., 2020). We used a RITE-cassette that allows the switch from a GFP-labelled protein to a new protein that is not fluorescently labelled (RITE(GFP-to-dark)) (Onischenko et al., 2020, Kralt et al., 2022). When tagging a Nup that stably associates with the NPC, we expected that successive cell divisions would dilute labelled NPCs by inheritance to both mother and daughter cells leading to a reduced density of labelled NPCs. By contrast, a labelled dynamic Nup is expected to redistribute to all NPCs of the nucleus and would not remain with individual NPCs (Figure 1C&D). We term this approach **d**ilution **o**f **RITE**-labelled complexes (doRITE). As a proof of concept, we performed doRITE on Nups previously reported to exhibit either stable (Nup133, Nup188) or dynamic (Nup60) association with the NPC (Rabut et al., 2004; Onischenko et al., 2020; Hakhverdyan et al., 2021). Upon addition of β-estradiol to induce Cre recombinase, >90 % of the cell population performed the genetic switch in less than one hour (Supplemental Figure 1A). We imaged cells at different time points after the induction of recombination while the cells were growing exponentially (Figure 1E). As the cells divided, different patterns were observed for the stable and dynamic Nups. For the stable Nups, Nup133 and Nup188, foci likely from individual NPCs could be clearly distinguished after 7 hours and their intensity was stable until only a few foci remained per cell. In contrast, the signal of Nup60 gradually decreased on all NPCs until it became undetectable over background at 11 hours (Figure 1E). Since the degradation rate of Nup60 is similar to other Nups (Hakhverdyan et al., 2021), reduction of its signal in doRITE is unlikely to be caused by protein degradation. doRITE can thus distinguish between stable and dynamically associated Nups.

To determine whether the fluorescent foci observed in doRITE at late timepoints correspond to individual NPCs, we analysed their intensity and inheritance patterns. Nup133-GFP foci exhibited a narrow intensity distribution indicative of a single particle species and the intensity remained stable after single foci could be distinguished (radial intensity profiles in Figure 1E and Supplemental Figure S1B). The foci did not cluster at the spindle pole body, as has been described for certain compromised NPCs (Heath et al., 1995, Supplemental Figure S1C). Furthermore, 43 ± 2% (mean ± s.d.) of the foci were inherited by daughter cells (Figure 1F), which is similar to the inheritance rate previously reported for pre-existing NPC (38 ± 12%, Khmelinskii et al., 2010). The average number of Nup133-RITE(GFP-to-dark) and Nup188-RITE(GFP-to-dark) foci per cell decayed exponentially (Figure 1G). By fitting this decay using the experimentally determined growth rate of the cells, we estimate an average of 158 ±22 (95% confidence interval) NPCs per cell at the time of recombination induction. This is at the upper end of the range expected in yeast cells (Winey et al., 1997), but the analysis likely overestimates the true number of NPCs because labelled Nups present in the soluble pool at the time of recombination continue to be incorporated and because we consider the induction of RITE as the timepoint 0 even though recombination occurs asynchronously and with some delay in the cell population (Supplemental Figure 1A).

In summary, our data are consistent with the interpretation that the observed foci represent individual NPCs. We conclude that doRITE can be used (1) to analyse the association dynamics of Nups at individual NPCs *in vivo*, (2) to follow the inheritance of individual NPCs from mother to daughter cell, and (3) to obtain single-NPC resolution in live budding yeast cells with standard diffraction-limited microscopy.

### Mlp1/2 and Pml39 associate with the NPC in a stable manner

Based on fluorescence-recovery-after-photobleaching (FRAP) experiments, it was previously reported that the nuclear basket Nups Mlp1 and Mlp2 exchange at the NPC within seconds (Niepel et al., 2013). However, a recent metabolic labelling-based biochemical analysis that systematically determined the exchange rate of all Nups on NPCs found the residence half-lives of Mlp1 and Mlp2 to be 4-5 hours (Hakhverdyan et al., 2021). To determine whether the basket filaments form stable or dynamic assemblies, we performed doRITE with Mlp1 and Mlp2 to visualize their association dynamics with NPCs in living cells. For comparison, we also performed doRITE with a set of additional Nups from different NPC subcomplexes, which exhibited different dynamic behaviours by metabolic labelling (Hakhverdyan et al., 2021) (Figure 2A). In this assay, Mlp1, Mlp2 and Pml39 behaved similarly to the stable Nups of group 1 (Nup133, Nup188, Pom152, Nup82, Nup159), which have an exchange half-life of greater than 7 hours at the NPC. Individual NPCs could be clearly distinguished starting 7 hours after induction of recombination, and the intensity of individual dots measured in radial intensity profiles remained stable through multiple cell divisions (Figure 2B, Supplemental Figure S2A). Similarly, Nup100 and Nup53, which biochemically behaved similarly to Mlp1 and Mlp2 in Hakhverdyan et al., 2021, remained stably bound at individual NPCs throughout our timecourse (Figure 2B, Supplemental Figure S2A). Thus, Nups of group 1 (residence time > 7 h) and group 2 (residence time 4.5-5 h) are stably bound at the NPC throughout multiple generations, although we cannot exclude the existence of an additional more dynamic pool of these proteins which could not be visualized by doRITE. In contrast, the analysed Nups of group 3, Nup116 (linker Nup) and Gle1 (cytoplasmic export platform), exhibited a gradual reduction of the GFP signal on all NPCs similar to Nup60 and consistent with a dynamic association with the NPC (Figure 2B and Supplemental Figure S2A).

**Figure 2.**
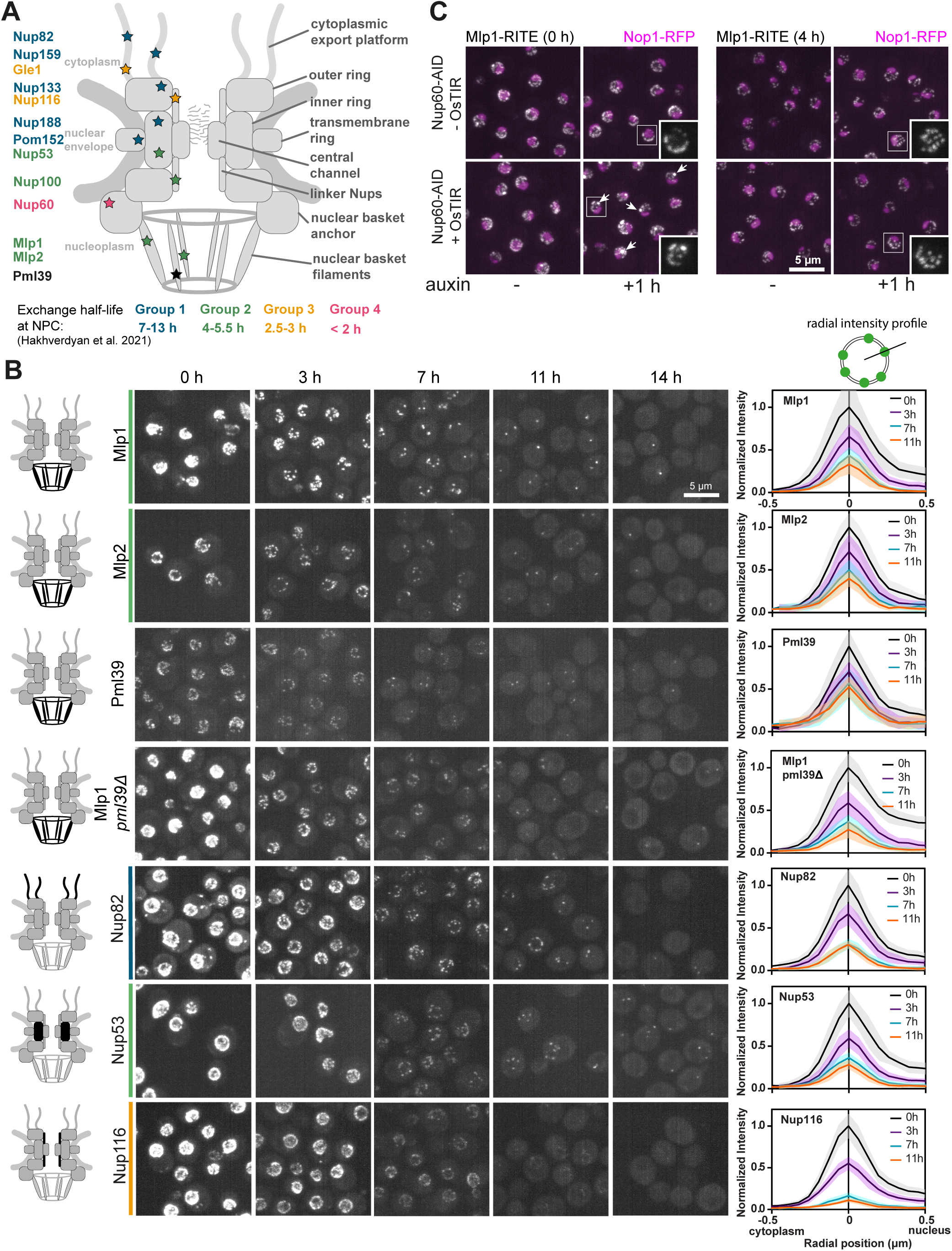
The Nups Mlp1, Mlp2 and Pml39 are stably associated with the NPC. (A) Overview of the Nups analysed. Approximate positions in the NPC are marked with stars. Colours correspond to the residence half-lives as characterized by Hakhverdyan et al., 2021. (B) Dilution of RITE(GFP-to-dark)- labelled Nups through multiple cell divisions. Cells expressing the indicated Nups endogenously tagged with the RITE cassette were imaged at different timepoints after induction of RITE. Maximum projections are shown. All images are adjusted to the same brightness and contrast values. Colour bars represent membership to dynamic group in Figure 2A. Schematics on left show location of Nup within the NPC. Graphs on right show radial intensity profile through fluorescent foci on the NE (compare schematic drawing on top) after indicated times of doRITE (Solid lines show mean of 20 cells, normalized for 0 h-timepoint, shaded area represents the standard deviation). (C) Mlp1-GFP remains at NPCs upon depletion of Nup60. Cells expressing Nup60 tagged with an auxin-inducible degron (AID) in the absence or presence of the E3 ligase adaptor OsTIR were treated with auxin for 1 h. Arrows indicate Mlp1-GFP foci formed by newly synthesized Mlp1-GFP which cannot incorporate into the NPC in the absence of Nup60. Maximum projections are shown. Nucleoli are labelled with Nop1-RFP (magenta).

The intensity of Mlp2-GFP and Pml39-GFP at the nuclear envelope and at individual foci was lower than that of Mlp1-GFP (Figure 2B), indicating that they are present at NPCs at a lower copy number. Quantification of the intensity of these GFP-tagged Nups on the NE in the non-nucleolar territory suggests that Mlp2 might be present in half (45.3±5 % (mean ± s.d.)) and Pml39 in one quarter (26.9±7 % (mean ± s.d.)) of the copy number of Mlp1 (Supplemental Figure S2B).

Pml39 anchors a subpopulation of Mlp1 to the NPC (Gunkel et al., 2023). To test whether Pml39 is required for the assembly of stable nuclear basket filaments, we performed doRITE with Nup133 and Mlp1 in *pml39Δ* cells. Similar to previous results (Gunkel et al., 2023) we observed an intensity reduction of Mlp1-GFP at the NE to 61±7% (mean ± s.d.) upon deletion of *PML39* (Supplemental Figure S2B). Mlp1-RITE(GFP-to-dark) still formed the characteristic NPC foci after 14 hours of doRITE in the absence of Pml39, but their intensity was reduced compared to wild type cells (Figure 2B, compare rows 1 and 4). This is consistent with the presence of a Pml39-dependent subpopulation of Mlp1 at the 14h-old NPCs, but indicates that Pml39 is not necessary for the stability of Mlp1 binding. Deletion of *PML39* or other nuclear basket components also had no effect on the dynamics of Nup133(GFP-to-dark) in doRITE (Supplemental Figure S2C).

Although Nup60 is essential for the recruitment of Mlp1/2 to the NPC, it was recently shown to be dispensable for the maintenance of already assembled Mlp1/2 at the NPC in meiotic cells (King et al., 2023). We tested whether this is also the case in exponentially growing cells. Indeed, we observed that depletion of Nup60 via an auxin-inducible degron tag (Nishimura et al., 2009) (Supplemental Figure S3A) did not affect the localization of Mlp1-RITE(GFP-to-dark) four hours after induction of RITE, suggesting that the old Mlp1 protein pool was maintained at NPCs (Figure 2C). In contrast, when all Mlp1 was labelled (no induction of RITE), a fraction of Mlp1-GFP accumulated in a nuclear focus reminiscent of the Mlp1-GFP focus formed in *nup60Δ* cells (Feuerbach et al., 2002; Niepel et al., 2013) (Figure 2C). Since this focus is not visible when only the old protein is labelled, we conclude that it is composed of newly synthesized Mlp1 that cannot be incorporated into the NPC in the absence of Nup60. Thus, Nup60 is required for the assembly of Mlp1 into the NPC but not for its maintenance. This appears to be evolutionarily conserved, as the same dependence was reported for the human homologues Nup153 and TPR (Hase & Cordes, 2003; Aksenova et al., 2020).

### Exclusion of old NPCs from the nucleolar territory is mediated by Mlp1

Next, we wanted to better understand the exclusion of Mlp1/2-positive NPCs from the nucleolar territory. Since integration of Mlp1 and Mlp2 into the NPC occurs on average within one hour of assembly (Onischenko et al., 2020), we expected that the NPCs visualized as individual foci by doRITE at the 14-hour timepoint would all carry Mlp1/2 and thus be excluded from the nucleolar territory. Indeed, only 10±3 % (mean ± s.d.) of 14 h-old Nup133-RITE(GFP-to-dark) NPCs were detected in the nucleolar territory marked by a nucleolar protein tagged with red fluorescent protein (Nop1-RFP) (Figure 3A). This finding is consistent with the observation that Mlp1-positive NPCs can occasionally localize to the nucleolus (Bensidoun et al., 2022). Deletion of *MLP2* did not significantly change this distribution, but in *mlp1Δ* and *mlp1Δ mlp2Δ* cells, the fraction of 14-hour old NPCs detected in the nucleolar territory increased to 15±2 % and 19±4 %, respectively (Figure 3A).

**Figure 3.**
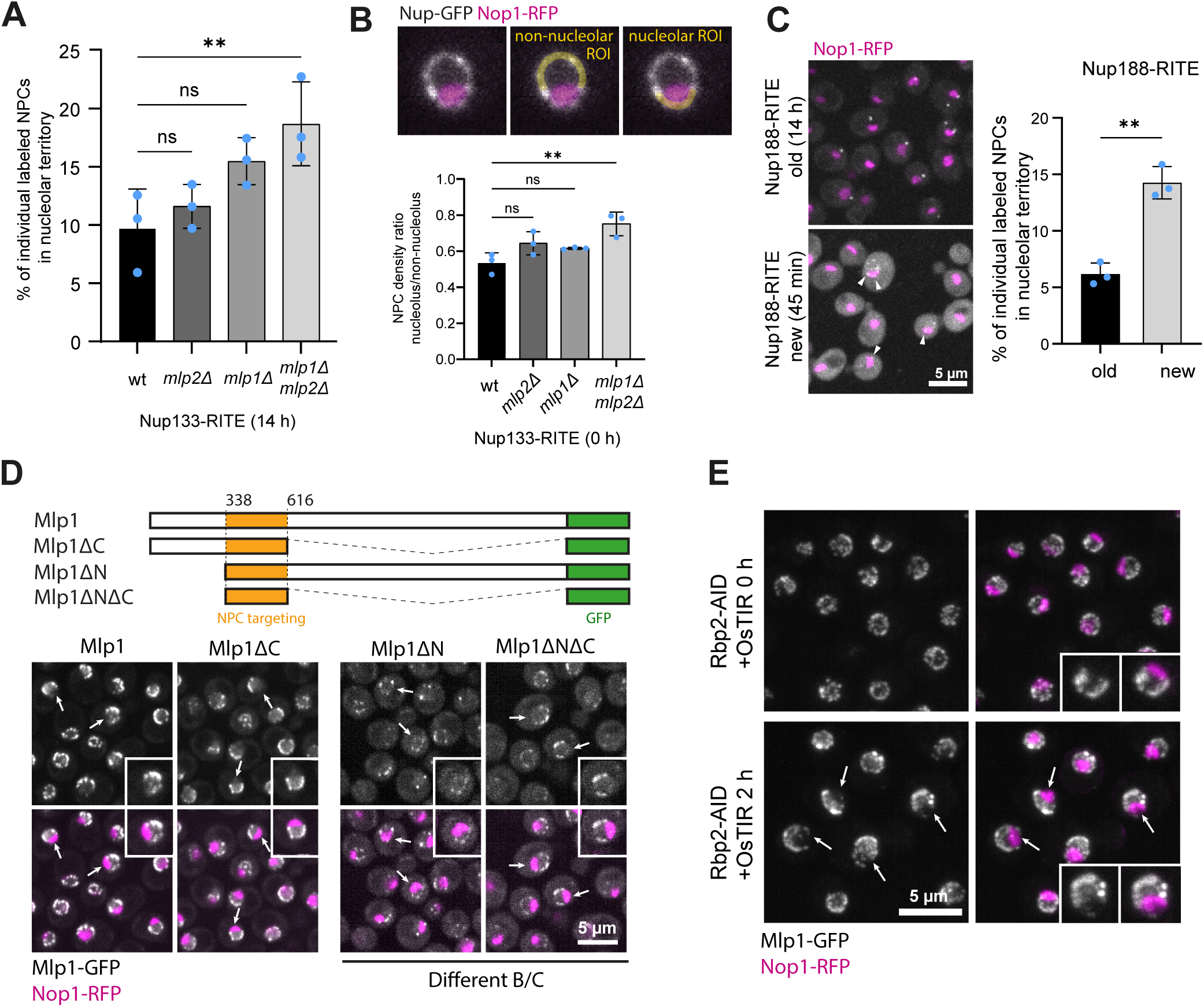
Exclusion of old NPCs from the nucleolar territory depends on Mlp1. (A) Percentage of Nup133-RITE(GFP-to-dark) labelled NPCs in the nucleolar territory in wild type cells and indicated mutants 14 h after RITE induction. At least 98 cells were analysed per condition and replicate by scoring the localization of NPCs in- or outside the nucleolar territory marked by Nop1-RFP. Blue dots show means of individual biological replicates and error bars the standard deviation. Stars indicate significance in one-way ANOVA with Sidak multiple comparison correction: ** p<0.005, ns not significant. (B) Top: Example of regions of interest (ROIs) used to quantify Nup-GFP intensity in the nucleolar and non-nucleolar region of the NE. Bottom: Mean ratio of Nup133-RITE(GFP-to-dark) in the nucleolar versus non-nucleolar territory without induction of recombination for wild type and nuclear basket deletion mutants. Blue dots show mean of biological replicates and error bars the standard deviation. At least 50 cells were analysed per replicate and condition. Stars indicate significance in one-way ANOVA with Sidak multiple comparison correction: ** p<0.005, ns not significant. (C) Percentage of GFP labelled NPCs in the nucleolar territory for Nup188-RITE(GFP-to-dark) (old) 14 h after induction of RITE and Nup188-RITE(dark-to-GFP) (new) 45 min after induction of RITE. Image panels show maximum projections. Arrowheads point to new NPCs in the nucleolar territory. At least 70 cells were analysed per strain and biological replicate by scoring the localization of NPCs in- or outside the nucleolar territory marked by Nop1-RFP. Blue dots show means of individual biological replicates and error bars the standard deviation. Stars indicate significance in unpaired t-test: ** p<0.005. (D) Mlp1 truncations (top) and their localization upon expression in cells (bottom). The truncations were introduced into a *mlp1Δ* background. All truncations are depleted from the nucleolar territory (arrows). Due to low expression of the constructs lacking the N-terminus, the right panels are adjusted with different brightness and contrast settings (Different B/C). The nucleolus is marked with Nop1-RFP. Maximum projections are shown. (E) Mlp1-positive NPCs remain excluded from the nucleolus upon depletion of PolII. RNA Polymerase II subunit Rbp2 was depleted from cells expressing Mlp1-GFP and Nop1-RFP via the auxin inducible degron. Depleted cells are larger and show increased nucleoplasmic signal of Mlp1-GFP. Arrows mark nucleoli with clear exclusion of Mlp1-GFP signal. Maximum projections are shown.

Not only are Mlp-positive NPCs present at a lower density in the nucleolar territory, but also overall NPC density is also reduced in this region (Wang et al., 2016). When we quantified the overall NPC density by measuring the GFP intensity of Nup133-GFP without the induction of RITE in the nucleolar and non-nucleolar sections of the NE, we also observed an increase in the nucleolar NPC density from 53±6 % in wildtype cells to 75±7% in the absence of Mlp1 and Mlp2 (Figure 3B). Importantly, doRITE with Nup133-RITE(GFP-to-dark) showed that the deletion of the basket protein genes did not compromise the stability of the NPC core (Supplemental Figure S3A). Thus, Mlp1/2 are indeed important for the exclusion of NPCs from the nucleolar territory.

Since newly assembled NPCs also lack Mlp1 and Mlp2, we tested whether they exhibit a similar distribution to 14 h-old NPCs in *mlp1Δ mlp2Δ* cells. We used an inverted RITE cassette (dark-to-GFP) to visualize newly assembled NPCs (Supplemental Figure S4A). Nup133-RITE(dark-to-GFP) generated a more diffuse NE signal at early timepoints that made the identification of individual newly assembled NPCs difficult – possibly due to the interaction of Nup133 with the membrane via an amphipathic helix (Nordeen et al., 2020). We therefore used Nup188 to compare the fraction of old (GFP-to-dark at 14 h) and new (dark-to-GFP at 45 min) NPCs in the nucleolar territory. We detected 6.1±1.0 % of 14-hour old NPCs in the nucleolar territory. In contrast, 14.2 ±1.4 % of newly assembled NPCs localized there (Figure 3C), which is comparable to the fraction of old NPCs observed in the nucleolar territory in *mlp1Δ mlp2Δ* cells. This suggests that in the absence of Mlp1/2, 14 h-old NPCs behave similar to newly assembled NPCs with respect to their nucleolar localization. In contrast, no difference in localization to the nucleolus was observed for old and new NPCs labelled with Mlp1-RITE (Supplemental Figure S4B). Since this population of “new” NPCs has already acquired Mlp1, this is consistent with the conclusion that the binding of Mlp1/2 leads to nucleolar exclusion of older NPCs.

Next, we tested which domain of Mlp1 mediates the exclusion from the nucleolus. A domain encompassing amino acids 338-616 of Mlp1 mediates localization to the NPC via binding to Nup60 and was reported to localize homogeneously to all NPCs (Niepel et al., 2013). We expressed truncated forms of Mlp1 tagged with GFP in *mlp1Δ* cells: the NPC binding fragment alone (ΔNΔC) or including the endogenous N- or C-terminal part of Mlp1 (Schematic representation Figure 3D). All three fragments were excluded from the nucleolar territory (Figure 3D, Supplemental Figure S5A), suggesting that the NPC-binding domain is sufficient to mediate the exclusion. Since we observed lower NE signal for Mlp1(ΔN)-GFP and Mlp1(ΔNΔC)-GFP, we expressed Mlp1(ΔNΔC)-GFP from a CEN plasmid in *mlp1Δ mlp2Δ* cells. In cells with low expression, Mlp1(ΔNΔC)-GFP was excluded from the nucleolar territory whereas in rare, highly expressing cells the construct localized homogenously to all NPCs, as previously reported (Niepel et al., 2013, Supplemental Figure S5B). This could indicate that the fragment preferentially binds NPCs in the non-nucleolar territory but can in principle also associate with NPCs the nucleolar territory.

The exclusion of Mlp1-positive NPCs from the nucleolar territory could be due to their retention in the non-nucleolar territory, e.g. via their interactions with nascent mRNA and mRNA maturation factors (Bermejo et al., 2011; Saroufim et al., 2015; Bensidoun et al., 2022). To test whether transcription affects nucleolar exclusion, we analysed Mlp1-GFP labelled NPCs in cells with an auxin-inducible degron tag on the Rpb2 subunit of Polymerase II (Pol II) (Derrer et al., 2019) (Supplemental Figure S5C). As reported in Bensidoun et al. (2022), we observed an increase in the nucleoplasmic signal of Mlp1 upon depletion of Pol II (Figure 3E). However, Mlp1-containing NPCs remained excluded from the nucleolar territory (Figure 3E). Thus, although inhibition of transcription affects nuclear basket assembly, the exclusion of Mlp1-containing NPCs from the nucleolus is maintained.

### Nup1 and Nup2 restrict the localization of NPCs to the nucleolar territory independent of Mlp1/2

Even in *mlp1Δ mlp2Δ* cells, the NPC density in the nucleolar territory remained reduced relative to the rest of the nuclear envelope (Figure 3B). This suggests that an additional, Mlp-independent mechanism is involved in restricting the localization of NPCs in the nucleolar territory. To test whether the nuclear basket contributes to this, we quantified the NPC density in the nucleolar and non-nucleolar territory in additional basket mutants.

Since *NUP1* is essential in our strain background, we used an auxin-inducible degron to deplete Nup1. After 2 hours of auxin treatment, when the protein is largely depleted (Supplemental Figure S6A), we observed a mild increase in the fraction of NPCs in the nucleolar territory (Figure 4A). Nup1 is not thought to be involved in the recruitment of Mlp1/2 to the NPC, and indeed Mlp1-localization upon depletion of Nup1 remained unchanged compared to the wild type and Mlp-positive NPCs remained excluded from the nucleolar territory (Figure 4B).

**Figure 4.**
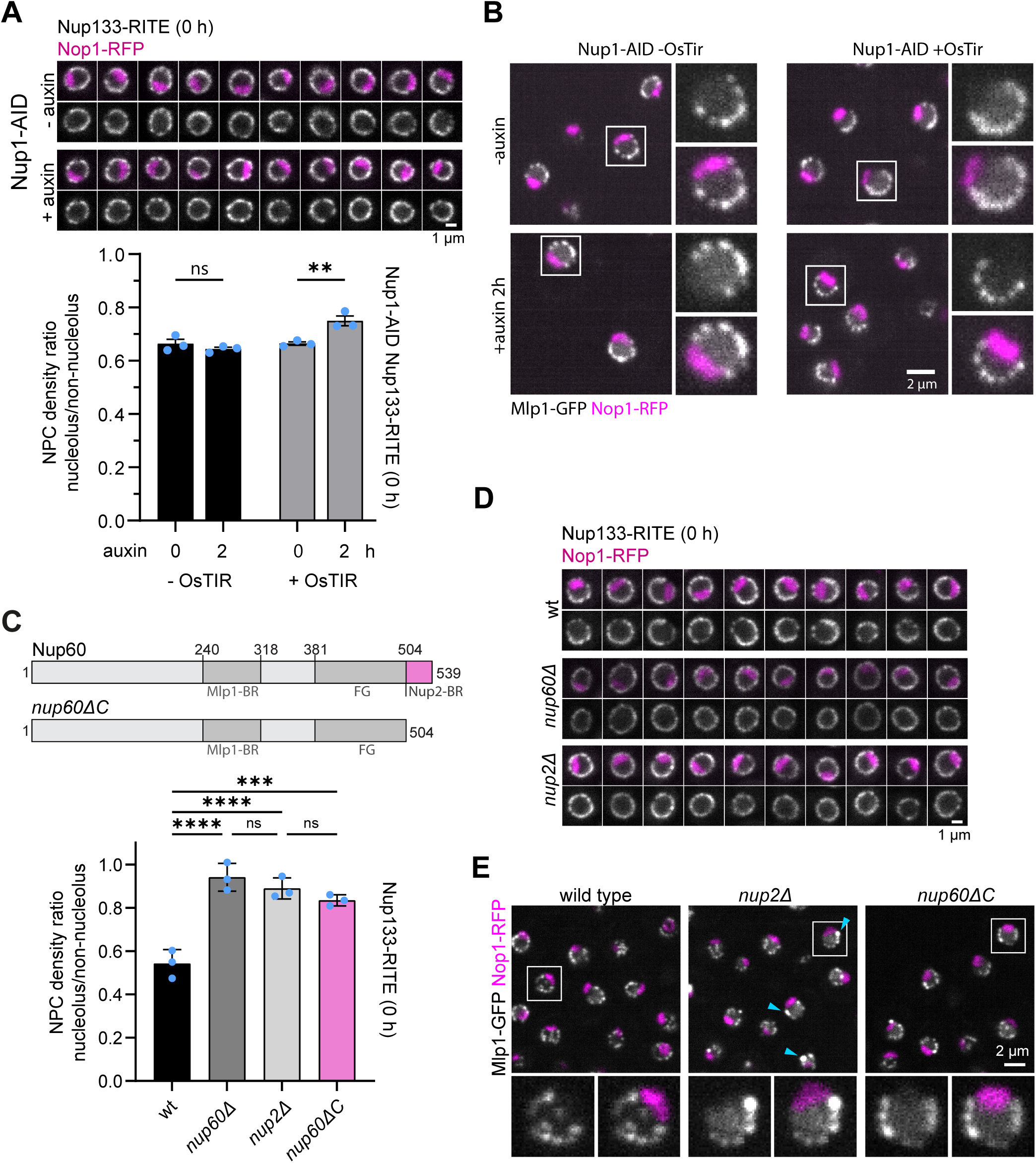
Exclusion of Mlp-negative NPCs from the nucleolar territory is mediated by Nup1 and Nup2. (A) Top: Representative nuclei of cells expressing Nup1 tagged with an auxin inducible degron (AID) before or after treatment with auxin for 2 hours. Cells express Nup133-RITE(GFP-to-dark) and Nop1-RFP and were imaged without induction of recombination. Single plane images are shown. Bottom: Mean NPC density in the nucleolar versus non-nucleolar territory of the strains shown above. (B) Cells expressing Mlp1-GFP and Nop1-RFP in cells expressing Nup1 tagged with an auxin inducible degron (AID) before or after treatment with auxin for 2 hours. Depletion occurs only in cells that also express OsTIR. Crops highlight exclusion of Mlp-positive NPCs from the nucleolar territory. Single plane images are shown. (C) Top: Scheme of Nup60 truncations. Mlp1-BR: Mlp1 binding region; Nup2-BR: Nup2 binding region. Numbers represent amino acid positions (drawn to scale). Bottom: Mean ratio of Nup133-RITE(GFP-to-dark) in the nucleolar versus non-nucleolar territory without induction of recombination for the indicated mutants. (D) Representative nuclei of wild type*, nup60Δ* and *nup2Δ* cells expressing Nup133-RITE(GFP-to-dark) and Nop1-RFP. Cells were imaged without induction of recombination. Single plane images are shown. (E) Cells expressing Mlp1-GFP and Nop1-RFP in wildtype and the indicated mutant backgrounds. Cyan arrows indicate Mlp1-GFP foci in nup2*Δ* cells. Crops highlight exclusion of Mlp-positive NPCs from the nucleolar territory. Single plane images are shown. All bar graphs: Blue dots show mean of biological replicates and error bars the standard error of mean. At least 50 cells were analysed per replicate and condition. Star indicates significance in one-way ANOVA with Sidak multiple comparison correction: **** p<0.0001, *** p<0.005, ** p<0.05, ns not significant.

Next, we tested the effect of deleting *NUP60*. This lead to a nearly homogenous distribution of NPCs around the nuclear envelope (Figure 4C&D). To distinguish effects of *nup60Δ* mediated through its role in recruiting Mlp1/2 from effects independent of Mlp1/2, we also determined the NPC density in the nucleolar territory upon acute depletion of Nup60 via auxin induced degradation (Supplemental Figure S6B, C). As we show in Figure 2C, Mlp1 is maintained at the NPC in this condition and Mlp-positive NPCs remain excluded from the nucleolar territory. However, overall nucleolar NPC density increased (Supplemental Figure S6C), suggesting that the increase in nucleolar NPC density in *nup60Δ* was at least in part independent of Mlp1/2 and likely due to the increased presence of Mlp-negative NPCs in this region of the nuclear envelope.

Nup60 also recruits Nup2 to the NPC via its C-terminus (Dilworth et al., 2001; Denning et al., 2001; Cibulka et al., 2022), Supplemental Figure S6D). Both, deletion of *NUP2* and deletion of the C-terminus of Nup60 (*nup60ΔC*) increased the NPC density in the nucleolar territory to similar levels and nearly as much as deletion of *NUP60* alone (Figure 4C), suggesting that the effect of *nup60Δ* is mediated predominantly through Nup2.

Again, we asked whether this effect was related to changes in the localization of Mlp-positive NPCs. We observed some mislocalization of Mlp1-GFP into foci in *nup2Δ* cells (Figure 4E, arrowheads, and quantification Supplemental Figure S6E). In *nup60ΔC* cells, Mlp1-GFP foci were much less frequent and smaller. Since intranuclear Mlp1 foci also form upon overexpression of Mlp1 (Strambio-de-Castillia et al., 1999; Kosova et al., 2000), we checked Mlp1 expression levels in these strains, but Mlp1-GFP expression was unchanged (Supplemental Figure S6F). In both strains, the Mlp1-GFP signal at the nuclear envelope appeared similar to wildtype cells, and Mlp1-GFP labelled NPCs remained excluded from the nucleolar territory (Figure 4E). Thus, Nup2 plays an Mlp-independent role in reducing NPC density in the nucleolar territory.

Taken together, we conclude that at least two independent molecular mechanisms lead to a reduced NPC density in the nucleolar territory. First, NPCs become depleted from the nucleolar territory once they bind the nuclear basket proteins Mlp1/2. Second, Nup2 and Nup1 mediate the indiscriminate exclusion of additional NPCs from the nucleolar territory. At present it is unclear whether the activities of Nup1 and Nup2 are independent of each other or contribute to the same pathway.

### Basket components affect the mobility of NPCs in the NE

The partial depletion of NPCs from the nucleolar territory could be caused by interactions that prolong the time spent in the non-nucleolar territory, since the nuclear basket is known to interact with multiple partners that can bind to chromatin (reviewed in Sumner & Brickner, 2022). More interactions in the non-nucleolar territory should manifest in a lower mobility of the NPCs in this region in wild type cells. In contrast, in *nup60Δ* cells, where we observe a homogeneous distribution of NPCs (Figure 4C&D), the mobility of NPCs would be expected to be the same throughout the NE. To test this idea, we applied simulations and tracking of individual NPCs using our probabilistic method, ExTrack (Simon et al., 2023).

First, we confirmed in a 2D simulation that a local decrease in the diffusion coefficient indeed causes a local increase in particle density. Particles diffusing in a 2D plane accumulated with higher density in the region with the lower diffusion coefficient (Figure 5A). Next, we compared the diffusion of Nup133-RITE(GFP-to-dark)-labelled NPCs 14 h after induction of recombination in wild type cells (Movie 1, Figure 5B) and in different nuclear basket deletion mutants (*nup60Δ*, *mlp1Δ*, *mlp2Δ* and *mlp1Δ mlp2Δ)*. We reasoned that these 14 h-old NPCs, which would be predominantly Mlp-positive in wild type cells and Mlp-negative in basket mutants, should show differences in mobility, if mobility is influenced by the presence of Mlp1/2. In wild type cells, NPC mobility exhibited behaviour switching between immobile, slow diffusive, and fast diffusive states (states 1, 2, & 3, Figure 5C&D). The diffusion coefficients of these states were similar in all analysed strains (Supplemental Figure S7A), but the fractions of NPCs that occupied the different states was changed in the mutants: NPCs in *mlp1Δ mlp2Δ* double mutants and *nup60Δ* mutants spend more time in the fast diffusive state relative to the wild type (Figure 5D). This shift in distribution translates into an overall increase in the average diffusion coefficients in *mlp1Δ mlp2Δ* and *nup60Δ* strains to 63±8 % and 53 ±14 % higher than in the wild type, respectively (Figure 5E). This is consistent with previous FRAP experiments that showed increased mobility of NPCs in *mlp1Δ mlp2Δ* cells (Niepel et al., 2013; Spichal et al., 2016). To test if this change in the movement behaviour could explain the changes in NPC distribution, we modelled the diffusion of NPCs along a spherical nucleus, applying diffusion parameters from *nup60Δ* cells in a cap region representing the nucleolar territory and from wild type cells in the rest of the NE (Figure 5F, see Track Simulations in the Methods). This relatively simple simulation generates a model where at steady state, the density of NPCs in the nucleolar territory would be 76 ±3 % of that in the non-nucleolar territory (Figure 5G).

**Figure 5.**
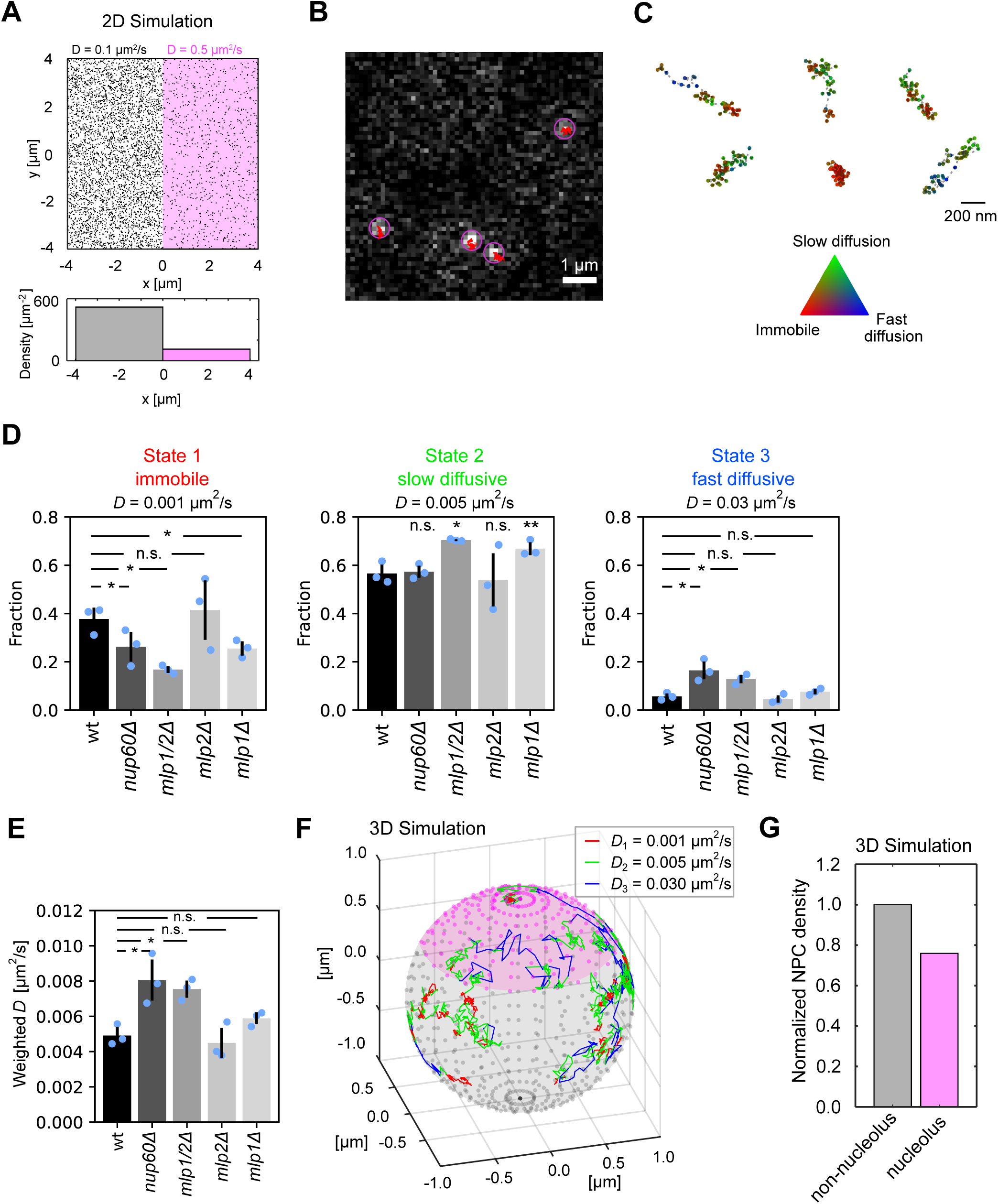
The nuclear basket proteins influence the mobility of NPCs. (A) 2D simulation of 1000 particles diffusing for 2000 timesteps in two territories with different diffusion coefficients as indicated (positions in the final time point are shown). The bar graph in the bottom shows the densities of the particles found in each area at the end of the simulation. (B) Still from Movie 1 showing four NPC-foci detected by Trackmate (magenta circles) with the corresponding track superimposed in red. (C) Example NPC trajectories from wild type cells. The ternary colour plot shows the probability of being in a specific state at each time-point. (D) Fraction of particles in different diffusion states for strains carrying deletions for different nuclear basket components. Tracks of NPCs labelled with Nup133-RITE(GFP-to-dark) obtained 14 hours after RITE induction were analysed by ExTrack with a 3-state model (immobile, slow diffusive and fast diffusive) with fixed diffusion coefficients as indicated. Bar graphs display the fraction of particles in each state computed from the transition rates between states. Results from individual biological replicates are shown as blue dots, the error bars represent the standard deviation of the three replicates. Stars indicate significance in pairwise t-tests comparing each mutant to the wild type: * p<0.05, ** p<0.01, ns not significant. At least 1100 tracks were analysed per condition and replicate. (E) Average diffusion coefficients derived from data displayed in (D) by weighting the three diffusion coefficients according to their respective fractions. The error bars represent the standard deviation. Stars indicate significance in pairwise t-tests comparing each mutant to the wild type: * p<0.05, ns not significant. (F) Simulation of particles diffusing on a sphere with three movement states. In the grey area, which represents the non-nucleolar territory, particles transition between states according to the rates extracted from tracking data in the wild type cells, while transition rates in the pink area, which represent the nucleolar territory, correspond to the ones from *nup60Δ* cells (Figure S7B). Three example tracks are shown and coloured according to the state the particle occupies in each timestep. (G) Steady state particle density in the nucleolar region relative to the density in the non-nucleolar regions obtained from the simulation model shown in (F). Data is based on 1000 tracks with 2000 timesteps each and normalized to the density in the non-nucleolar territory.

Our data and simulation thus support the hypothesis that an increased apparent diffusion coefficient of NPCs on the nuclear envelope contributes to the reduced NPC density in the nucleolar territory (Figure 5A, F&G). Indeed, the tracks from *mlp1Δ mlp2Δ* and *nup60Δ* cells exhibit a reduced frequency of binding events, that is, less transitions from a diffusive to a bound state (Supplemental Figure S7 B). This would be consistent with reduced interactions of NPCs with nuclear components in these strains. However, since the two mutants exhibit similar average diffusion coefficients (Figure 5E), the observed difference in mobility can probably be attributed to the loss of Mlp1/2 only, which is absent in both mutants. This interpretation is corroborated by the observation that acute depletion of Nup60, which does not affect the localization of Mlp1, did not affect NPC mobility (Supplemental Figure S7C). Thus, we cannot at this point separate the Mlp1-dependent and Mlp1-independent roles of Nup60. Furthermore, the nucleolar NPC density predicted in our model is still significantly higher than what we observe experimentally (76 ±3 % in the model versus 53±6 % in wild type cells, compare Figure 5G to Figure 4B). Therefore, while our results support the hypothesis that NPC distribution can be the result of differential mobility caused by different interaction networks in the different regions of the nucleus, additional mechanisms are likely to contribute (Figure 6).

**Figure 6:**
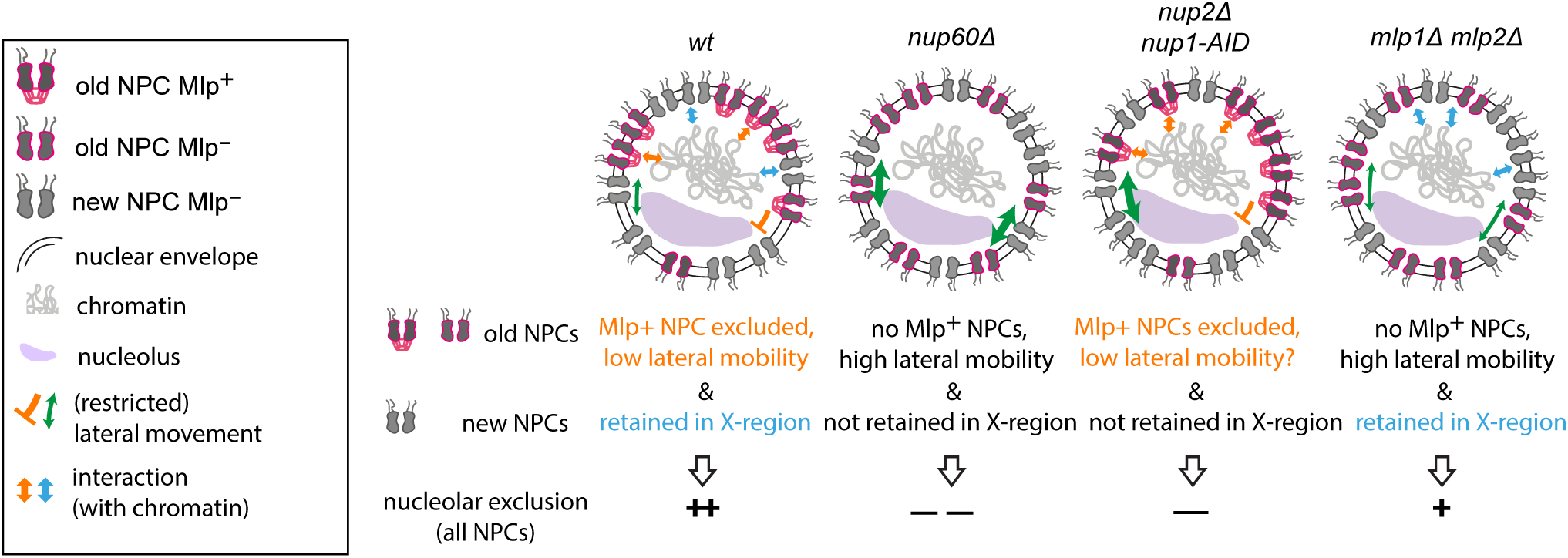
Model of how nuclear basket proteins regulate the distribution and movement of NPCs on the nucleus. Interactions with the nuclear interior (blue arrows) and lateral diffusion along the nuclear envelope (green arrows) govern the distribution of NPCs on the nuclear surface. Mlp1/2 and Nup1/2 independently contribute to reduced NPC density in the nucleolar territory.

## Discussion

### doRITE as a versatile tool for single NPC analysis in yeast

To understand the differences between NPC variants, it is crucial to develop methods that do not only report on bulk behaviour but can distinguish between subpopulations of NPCs and ideally provide single NPC resolution. While single NPCs can readily be resolved by confocal microscopy on the flat bottom of many adherent mammalian cell types, this poses a greater challenge in budding yeast, which have small, spherical nuclei with high NPC density. In this work, we developed RITE into a versatile tool to achieve single NPC resolution in *S. cerevisiae* by utilizing the properties of yeast NPCs: (i) high stability of the complex, (ii) multiple copies of the labelled protein per complex, and (iii) intact inheritance through multiple cell cycles (Figure 1D&E). We harnessed the power of this method to analyse the exchange dynamics of different Nups at the NPC (Figure 2), to determine the localization of single NPCs relative to nuclear landmarks (Figures S1B, 3&4) and to track the movement of NPCs along the NE (Figure 5). The NPC is particularly suitable for doRITE due to the presence of several copies of the labelled protein per complex which yields a high signal-to-noise ratio and enables tracking of individual NPCs. However, doRITE could in principle also be applied to other stable complexes such as the ribosome.

### The nuclear basket filaments form a stable structure on the NPC

FRAP is a common method to investigate the dynamics of biological molecules and has been used to systematically characterize Nup dynamics in mammalian cells (Rabut et al., 2004). However, only highly dynamic interactions at NPCs, like those of transport receptors, can be quantified by FRAP in budding yeast (Derrer et al., 2019), since longer measurements are confounded by the mobility of entire NPCs on the nuclear surface. doRITE overcomes this limitation and provides a complementation to biochemical assays, which were recently used to determine the exchange rate of Nups on the NPC in bulk (Onischenko et al., 2020; Hakhverdyan et al., 2021).

Our results are largely in agreement with the previously reported overall categorisation of Nups as stable or dynamic based on biochemical data (Figure 2, (Hakhverdyan et al., 2021)). However, we observed that the Nups with reported residence times at the NPC of 4.5-5 hours (Mlp1, Mlp2, Nup53 and Nup100) exhibit stable properties in our doRITE assay, similar to the stable core Nups of group 1 with a residence time of > 7 hours (Figure 2B). This discrepancy could arise from the presence of two pools with different dynamic behaviour, of which the mass-spectrometry approach would observe the average while our doRITE approach would primarily visualize the more stable pool. Alternatively, ongoing assembly of Mlp1/2, which join the NPC only 40 min after the bulk of the other Nups (Onischenko et al. 2020), might contribute to a higher apparent exchange of these Nups during the chase period in Hakhverdyan et al., 2021.

The interpretation of Mlp1 as a stable Nup is corroborated by the observation that, once assembled, it is maintained at the NPC independent of Nup60 (Figure 2C), as also reported for meiotic budding yeast and mammalian cells (Aksenova et al., 2020; King et al., 2023). Furthermore, the human homologue TPR exhibits stable association with the NPC by FRAP (Souquet et al., 2018). Interestingly, Mlp1 also forms stable assemblies at the NPC when *PML39* is deleted (Figure 2B), suggesting that neither Pml39 itself nor the additional copies of Mlp1 recruited to the NPC by Pml39 (Gunkel et al., 2023) are required for the assembly of stable nuclear basket filaments.

Although we present clear evidence for the stable assembly of Mlp1, Mlp2 and Pml39 at the nucleoplasmic face of the NPC, we cannot exclude the possibility that a second, more mobile pool exists which might be associated with the NE more generally, as suggested by labelling at NE sites away from the NPCs by immuno-gold (Strambio-de-Castillia et al., 1999). Furthermore, transient interactions via Nup60 could prevail during early assembly stages before being converted into more stable interactions during maturation of the nuclear basket. Such maturation might be correlated with the assembly of the second nucleoplasmic Y-complex ring, which was observed in a fraction of yeast NPCs (Akey et al., 2022), since density for nuclear basket filaments was recently resolved on only those NPCs with two nucleoplasmic Y-complex rings by *in situ* cryo-electron tomography and tomogram averaging (Singh et al., 2024). Interactions with the Y-complex may also be responsible for maintaining Mlp1/2 at the nuclear basket upon depletion of Nup60 (Stankunas & Köhler, 2024; Singh et al., 2024) .

Quantification of the amounts of GFP-tagged basket components at the NE suggest that the relative abundance of Mlp1:Mlp2:Pml39 is 4:2:1 (Supplemental Figure S2B). The structure and stoichiometry of the majority of the core NPC is now well characterized thanks to high resolution cryo-electron tomography data and structural modelling (Kim et al., 2018; Allegretti et al., 2020; Akey et al., 2022, 2023), but although density for the nuclear basket was recently observed (Singh et al., 2024), it does not yet have sufficient resolution to determine the absolute copy number of all nuclear basket components. Based on the stable complex formed and its well-structured appearance as a “basket” (Kiseleva et al., 1998), we consider it likely that the different Nups are present at a fixed stoichiometry. Copy numbers of 16 and 8 for Mlp1 and Mlp2, respectively, would be in the range of previous mass-spectrometry-or fluorescence microscopy-based quantifications (Mi et al., 2015; Kim et al., 2018). Surprisingly, this would suggest a copy number as low as four for Pml39, which would be the first instance of a Nup which is present in less than one copy per spoke. Since Pml39 can interlink Mlp1 molecules (Gunkel et al., 2023), we propose that Pml39 might connect the basket proteins of neighbouring spokes. This would be consistent with the localization to the distal ring reported for the mammalian homologue ZC3HC1 (Gunkel et al., 2021).

### NPC retention in the non-nucleolar territory of the NE is mediated by the basket proteins

It has previously been observed that Mlp-positive NPCs are excluded from the nucleolar territory (Galy et al., 2004; Niepel et al., 2005). In this work, we show that this phenomenon affects predominantly old NPCs (Figure 3A-C), which is in agreement with the observation that Mlp1/2 are assembled into NPCs after a long delay. We consider two hypotheses for the molecular mechanism of exclusion: retention in the non-nucleolar region due to interaction with chromatin and/or mRNA (Figure 6, blue arrow), or a barrier that prevents entry of Mlp-positive NPCs into the nucleolus (Figure 6, red inhibition signal). Depleting RNA Pol II did not cause Mlp-positive NPCs to redistribute to the nucleolar region (Figure 3E), suggesting that exclusion does not rely on tethering to nascent mRNA or transcribing gene loci. Since even the minimal NPC binding domain of Mlp1 exhibited preference for the NPCs outside of the nucleolar territory (Figure 3D), nucleolar exclusion might also involve regulation through the Mlp1 binding site. Further studies are needed to determine exactly which molecular mechanisms are at play.

Alternatively, exclusion could be mediated by the specific biophysical properties of the nucleolus (reviewed in Tartakoff et al., 2022), which could present a barrier for entry to Mlp-positive NPCs. Since the free pool of Mlp1 in principle has access to the nucleolus (Bensidoun et al., 2022), it is possible that the barrier applies only to the NPC as a whole, or to the fully assembled basket.

In addition, we describe a second, Mlp-independent mechanism that leads to a reduced density of NPCs in the nucleolar territory and relies on Nup2 and to a lesser extent on Nup1 (Figure 4A, C-D). Nup2 and its homologues in *Aspergillus nidulans* and metazoa have been shown to interact with chromatin (Ishii et al., 2002; Markossian et al., 2015; Holzer et al., 2021), which could lead to the preferred localization of NPCs to the non-nucleolar region of the NE. Mlp1/2 independent interactions of NPCs with chromatin are also consistent with a recent study that demonstrated that Mlp-negative NPCs are involved in subtelomeric gene silencing (Choudhry et al., 2023). Alternatively or in addition, the FG repeats of Nup1 and Nup2, which are thought to extend into the nucleoplasm, might form a condensate that is non-miscible with the nucleolar condensate and therefore disfavours localization to the nucleolar territory. Either mechanism may apply also for the soluble form of Nup2, since it is clearly present at a strongly reduced density in the nucleolar territory (Supplemental Figure S6D).

### Tracking single NPCs in budding yeast

Using doRITE, we were able to track the diffusion of individual NPCs on the nuclear envelope of budding yeast up to 30 s, yet, many tracks were shorter, since NPCs diffused out of the focal plane. However, employing a probabilistic approach that considered the dynamics and populations in groups of single particle trajectories (Simon et al., 2023), we could determine the diffusion coefficients of NPCs in different mobility states from our tracking data. Our analysis overall supports the hypothesis that the modulation of diffusion due to differential interactions can lead to a non-homogeneous distribution of NPCs on the NE (Figure 5). In the future, the tracking and analysis pipeline we developed will be applicable to further study the forces that act on NPCs and drive their movement and distribution on the nucleus. Unfortunately, we could only implement single NPC tracking for old NPCs, since new NPCs labelled with an inverted RITE-tag did not have enough signal-to-noise ratio. As a result, we could not track young, Mlp-negative NPCs and could therefore not directly determine the effect of Nup2 or Nup1 on NPC mobility.

In conclusion, we have developed and applied a versatile new tool to better understand the properties and organization of the nuclear basket. We demonstrate that Mlp1/2 and Pml39 form a stable nuclear basket which does not depend on Nup60 for maintenance and show that multiple pathways contribute to regulating the distribution of NPCs on the nuclear envelope. Our results on nuclear basket dynamics and binding interdependencies are very consistent with findings in metazoa, indicating a high level of conservation in the building principles of the basket. However, the interplay between the nucleolus and NPCs we describe in yeast may be of limited relevance in metazoa, since nucleoli are not directly associated with the nuclear envelope in most metazoan cell types.

Furthermore, in contrast to yeast, metazoa undergo open mitosis during which NPCs disassemble, meaning that their age matches the age of the cell. NPCs in old, post-mitotic cells have been shown to exhibit signs of malfunction (D’Angelo & Hetzer, 2008), and it will be interesting to determine whether old NPCs in young yeast cells exhibit similar deficiencies.

## Materials and Methods

### Yeast strain and plasmid construction

*Saccharomyces cerevisiae* strains were created in the BY4742 wild type strain background using standard yeast genetic techniques by transformation of either a PCR product (Longtine et al., 1998) or a linearized plasmid with homology regions to the yeast genome. Some strains with multiple genetic modifications were generated by mating, sporulation and spore selection. Plasmids were generated using standard restriction-based cloning methods. The V5 tag in the original RITE plasmid pKV015 from (Verzijlbergen et al., 2009) was replaced due to the presence of unfavourable codons that led to low expression levels of tagged proteins in yeast. Used strains, plasmids and primers are listed in Supplemental Tables 1-3.

### Yeast culture conditions and treatments

Yeast cultures were grown to saturation and diluted to OD=0.01 in synthetic complete medium containing 2% glucose at 30°C. Prior to induction of recombination, cells were kept and grown on selective (-ura or hygromycin) solid or liquid medium to select against a low level of uninduced recombination from the Cre recombinase which is continuously expressed under control of a TDH3 promoter (Terweij et al., 2013). Exponentially growing cells were induced to recombine with 0.5 µM β -estradiol (Sigma), and cultures were imaged in exponential growth phase after the indicated times, if not mentioned otherwise. Auxin-inducible degradation of proteins was induced by treatment with 0.5 mM indole-3-acetic acid (IAA) (Sigma, I2886-5G, CAS: 87-51-4) and 4 µM phytic acid dipotassium salt (Sigma, 5681, CAS: 129832-03-7) for the indicated time.

### Microscopy

Cells were imaged in 384-well plates (Matrical) coated with concanavalin A for immobilization. 3D volume stacks of yeast cells with RITE-labelled NPCs were acquired on a temperature-controlled inverted Nipkow spinning disk microscope equipped with the Yokogawa Confocal Scanner Unit CSU-W1-T2 SoRa and a triggered Piezo z-stage (Mad City Labs Nano-Drive). It was used in spinning disk mode with a pinhole diameter of 50 µm combined with a 1.45 NA, 100x objective and controlled by the NIS Elements Software (Nikon). Images were acquired with a sCMOS Hamamatsu Orca Fusion BT camera (2304 x 2304 pixel, 6.5 x 6.5 μm pixel size). Imaging was performed at 30 °C with 50 % laser intensity of a Oxxius 488 nm 200 mW LNC Version and/or 5% laser intensity of Oxxius 561 nm 200 mW LNC Version light source. Z-stack data were taken with the Piezo stage in triggering mode with 0.2 µm sectioning. Exposure times were 200 ms for GFP and 50 ms for RFP.

NPC-tracking experiments were performed on a temperature-controlled inverted Nipkow spinning disk microscope equipped with the Yokogawa Confocal Scanner Unit CSU-W1-T2 controlled by the VisiVIEW Software (Visitron). It was used in spinning disk mode with a pinhole diameter of 50 µm combined with a 1.45 NA, 100x objective. The microscope is equipped with two EMCCD Andor iXon Ultra cameras (1024×1024 pixel, 13×13um pixel size) and was used in a dual camera mode for measuring distances between NPCs and spindle pole bodies, in single camera mode for all tracking experiments. Imaging was performed at 30 °C with 80 % laser intensity of a DPSS 488 nm 200 mW and/or Diode 561 nm, 200 mW light source. Timelapse data were taken with 100 ms exposure time in stream mode for 300 frames. Z-stack data for Supplemental Figure S1C were taken with the Piezo stage in triggering mode with 0.2 µm sectioning.

### Quantitative real-time PCR

For qPCR, cells at OD600 0.8–1 (2 ml) were harvested by centrifugation. DNA was extracted from cells disrupted with glass beads in lysis buffer (1 % SDS, 2% Triton-X, 100 mM NaCl, 10 mM Tris pH 7.4, 1mM EDTA) with Phenol:Choloform:Isoamyl alcohol followed by ethanol precipitation. qPCR was performed on a StepOnePlus Instrument (Invitrogen) using PowerUp^TM^ SYBR^TM^ Green Master Mix for qPCR (Thermo Fisher Scientific). All experiments were carried out in three technical replicates and three biological replicates. Data were analyzed using a standard curve and calculated relative to NUP133 ORF as endogenous control.

### Western blotting

Logarithmically growing cells were collected and lysed with 0.1 M NaOH for 15 min. Lysate was resuspended in SDS-loading buffer and 1 OD of cells were separated in 10 % acrylamide SDS-PAGE gels. Blotting of proteins was done with a Biorad semi-dry system in TBE buffer (pH=8.3) or 20% methanol/10mM CAPS buffer (Sigma-Aldrich, C2632-250G) at 25 V and 1 Amp followed by a Ponceau-S quick protein stain. V5 epitope, Hxk1 and Crm1 detection was performed with the following antibodies: mouse-anti-V5 antibody (AbD Serotec, MCA1360), goat-anti-mouse coupled to Alexa680 (Thermo Fisher, A-21057), rabbit-anti-Hxk1 (USBiological, 169073), rabbit-anti-Xpo1 (Zeitler & Weis, 2004) and goat-anti-rabbit coupled to IRDye800 (LI-COR Biosciences GmbH, P/N 926-32211).

### Inheritance tracking

Images were acquired on the Visitron Spinning Disk microscope (see above). Cells were imaged 14 hours after RITE induction with β-estradiol at 3 min intervals in the brightfield channel to follow individual division events and identify mother-daughter pairs. At the end of the time-lapse movie, a single 3D stack was acquired in the GFP channel, and 100 mother-daughter pairs were analysed per biological replicate.

### Intensity analysis of nucleolar versus non-nucleolar NE

The intensity was analysed in three biological replicates on at least 50 nuclei per condition and replicate. On maximum z-projections, nuclei in which the nucleolus presented a moon-shaped side view abutting the NE were selected manually for analysis. For each nucleus, the z-slice with the maximum intensity for the nucleolar channel (Nop1-RFP) was selected for the intensity measurement. A hand-drawn line of width 3 pixels following the NE as delineated by the Nup-GFP signal was created in FIJI first on the non-nucleolar side of the nucleus and then on the nucleolar side. Out-of-cell background was subtracted, and the ratio between the nucleolar and the non-nucleolar signal calculated.

### Image and Data analysis

Image analysis and adjustments were carried out in FIJI (Schindelin et al., 2012). Data analysis, curve fitting, statistical analysis and plotting was carried out in Excel, Matlab or GraphPad Prism. Analyzed cell numbers are listed in Supplemental Table 4 if not specified elsewhere. For line scans quantifying NE signal in doRITE (Figures 1E, 2B, S2A), 20 cells were picked on maximum projections, the central slice of the z-stack was determined, and a line drawn perpendicular to the plane of the NE through the brightest spot on the NE. The profiles were aligned at the peak intensity, averaged and normalized for each strain between background and peak intensity at the 0h timepoint.

### Single Particle Tracking

NPC trajectories were extracted from the movies using the FIJI (Schindelin et al., 2012) plugin Trackmate (Ershov et al., 2022). Briefly, peaks were identified using the LoG detector method with a radius of 300 nm and a quality threshold of 25. Tracks were then reconstructed from localizations using the option *simple LAP tracker* without gaps and a maximum linking distance of 300 nm. A mask based on thresholding and dilation was applied to exclude regions of high fluorescence intensity to avoid mislocalizations in regions with higher NPC densities. Additionally, tracks were filtered to have at least 5 time points, an average quality of at least 35, and a minimum quality of 25. To avoid tracks that might contain incorrectly linked localizations from one frame to the next, we also discarded tracks within 1 µm of one another. The resulting dataset was analysed using ExTrack, our probabilistic multi-state analysis pipeline (Simon et al., 2023). We used a 3-state model, since it fit the data much better than a 2-state model. For a data set of 57 tracks of 30 time points, the likelihood of the three-state model was 1.4×10^5^ higher than for two states (Vaart, 1998). We extracted the diffusion coefficients of the three states for each replicate of every condition (Supplemental Figure S7A). Due to the similarity across many tested conditions, subsequent analysis used the rounded average values for the three states, 0.001 µm^2^/s (immobile state), 0.005 µm^2^/s (slow state), and 0.03 µm^2^/s (fast state), to compare the transition rates between states (Supplemental Figure S7B). These estimated transition rates were used to compute the population fractions at equilibrium (Figure 5D). To calculate the average diffusion coefficients (Figure 5E), we computed the weighted average for each condition (*_Davg_ = F*_1_*D*_1_ + *F*_2_*D*_2_ + *F*_3_*D*_3_), where *F_N_* is the fraction in the Nth state and *D_N_* is the corresponding diffusion coefficient. Finally, for each individual track, we computed the probability of being in each state per timepoint (Figure 5C).

### Track simulations

To explain the differences of density between the non-nucleolar and nucleolar regions, we simulated tracks moving in an environment with two different diffusion characteristics in 2D (Figure 5A) and 3D (Figure 5F). In the 2D simulation, we characterized the impact of a change in diffusion coefficient on particle density by modelling two areas with distinct diffusion coefficients using 5000 trajectories with 5000 time steps each to find the distribution of positions at steady state. Figure 5A shows the final positions of each track. In the 3D simulations, we tested whether the observed differences in the nucleolar densities could be attributed to the differences of transition rates with the three motion states at fixed diffusion coefficients (0.001, 0.005 and 0.03 µm^2^/s. Tracks were simulated on a 1 µm radius sphere composed of a body and a circular cap whose surface area was one third of the total area to represent the nucleolar region. Figure 5F shows three example tracks. In these tracks, the states switched stochastically in time according to a Markov chain model of transition probability per step of p_12_ = 0.028, p_13_ = 0, p_21_ = 0.013, p_2_ = 0.041, p_31_ = 0, p_32_ = 0.15 in the cap (nucleolus) and p_12_ = 0.055, p_13_ = 0, p_21_ = 0.035, p_23_ = 0.02, p_31_ = 0, p_32_ = 0.2 in the body (non-nucleolus). These transition probabilities were chosen based on the measured values for the *nup60Δ* data representing the behaviour in the nucleolar territory, and the wild type data for the behaviour in the non-nucleolar territory. 1000 tracks of 2000 time steps were used to find the steady state, where the last position of each track was used to calculate the density.

To determine how the measured diffusion coefficients could be influenced by our 2D measurements of 3D diffusion, we repeated our 3D simulations for a single diffusion coefficient (Supplemental Figure S7D) and extracted the average diffusion coefficients as a function of axial position. Near the centre of the cell, that is where our experimental data is recorded, the 2D projection decreases the apparent diffusion coefficient by half (Supplemental Figure S7E). This indicates that our reported diffusion coefficients are likely an underestimate of the true values, but within a factor of two.

Notably, the differences of density depend on the ratios between diffusion coefficients. As these ratios are conserved in the 2D projections, our measured densities in simulations are not affected by the shift of diffusion coefficients.

#### Author Contributions

Conceptualization: JZ, LW and ED. Methodology: JZ, ED, FS, LW. Investigation: JZ, LI, GB, YK, AW and ED. Formal analysis: JZ, FS, LI, LW and ED. Simulations: FS, NK and LW. Writing – Original Draft: JZ, ED. Writing – Review & Editing: all authors, Funding acquisition: LW, ED. Supervision: JZ, LW, ED.

## Acknowledgements

We thank Alwin Köhler and Fred v. Leeuwen for providing plasmids, ScopeM and Joachim Hehl for microscopy support, Evgeny Onischenko, Stephanie Heinrich as well as Karsten Weis and members of his group for sharing resources, discussion and critical comments on the manuscript. The work was supported by an ETH Zurich research grant ETH-33 19-1 to ED, and in part with funding from the Polytechnique Montréal Direction de la recherche et de l’innovation (DRI) to NK and LW, the Canada First Research Excellence Fund (TransMedTech Institute) to NK and LW and the Natural Sciences and Engineering Research Council of Canada (NSERC Discovery grant, RGPIN-2022-05142) to LW. AW and YK acknowledge support from the Amgen Scholar Program.

## Legends Supplemental Figures

**Figure S1.** doRITE foci show the characteristics of individual NPCs. (A) Kinetics of genomic recombination upon induction of RITE. Quantitative real-time PCR was performed on genomic DNA extracted from Nup133-RITE(GFP-to-dark) cell populations harvested at different time points after RITE induction. Primers detected sequences present before (preRITE, filled circles) or after (postRITE, open circles) recombination. Amounts are shown relative to the NUP133 ORF, which is not affected by the RITE switch. preRITE amounts were normalized to timepoint before induction, postRITE amounts to the 24 hour timepoint. Datapoints represent means from three biological replicates, error bars represent the standard deviation. Red and blue lines are fits with a single exponential, half-times calculated from the fit are given in the legend. (B) Integrated and background subtracted intensity from individual NPC foci 8 or 14 h after RITE induction. 3D stacks were acquired and intensity was quantified on the best-in-focus slice. Top: Intensity distribution of 350 isolated foci. The red line represents a 2-component Gaussian fit. Bottom: Dots represent the centre peak position of the smaller component of the Gaussian fit for replicates of NPC intensity distributions 8 or 14 hours after RITE induction. Lines represent the median. Each distribution contains >200 individual foci. (C) NPC-foci do not cluster with SPBs. Distribution of distances of Nup133-RITE(GFP-to-dark) foci at 14 h after RITE induction from the spindle pole body (SPB) labelled by Spc42-mCherry. At least 500 SPB-NPC distances were measured per replicate. Blue dots represent fraction in individual biological replicates. Error bars represent standard error of the mean. Image shows a representative maximum projection of the analysed cells.

**Figure S2.** Nup dynamics determined by doRITE. (A) Dilution of RITE(GFP-to-dark)-labelled Nups through multiple cell divisions. Cells expressing the indicated Nups endogenously tagged with the RITE cassette were imaged at different timepoints after induction of RITE. Maximum projections are shown. All images are adjusted to the same brightness and contrast values. Cyan arrowhead highlights cytoplasmic Nup159 granule visible only before induction of RITE. Colour bars represent membership to dynamic group in Figure 2A. Schematics on left show location of Nup within the NPC. Graphs on right show radial intensity profile through fluorescent foci on the NE (compare schematic drawing on top) after indicated times of doRITE (solid lines show mean of 20 cells, normalized for 0 h-timepoint, shaded area represents the standard deviation). (B) The mean intensity of GFP-tagged nuclear basket proteins in the non-nucleolar NE region. Signal outside the nucleus was considered as background, but for the condition Mlp1 Δpml39 the mean signal between nucleoplasm and cytoplasm was considered as background due to higher nucleoplasmic Mlp1 signal. Background subtracted signals are normalized to Mlp1. Blue dots show mean of biological replicates of at least 41 nuclei analysed per condition and replicate. Schematic shows measured area (yellow) of the NE. (C) Mutations in basket Nups do not affect stability of the NPC scaffold. Cells expressing Nup133 endogenously tagged with the RITE cassette were imaged at different timepoints after induction of RITE. Maximum projections are shown. All images are adjusted to the same brightness and contrast values.

**Figure S3.** Depletion of Nup60 via auxin inducible degradation. Western blot of the depletion of Nup60, in the absence or presence of the E3 ligase adaptor OsTIR, by an auxin inducible degron (AID) at the indicated timepoints after addition of auxin (equivalent of 1 OD_600_ of cells was loaded in each lane). V5 antibody detects the Nup60-V5-AID as well as the RITE-tagged Mlp1. Crm1 was used as a loading control.

**Figure S4.** A RITE(dark-to-GFP) cassette to visualize newly assembled NPC. (A) Top: Schematic of the dark-to-GFP RITE cassette labelling newly forming NPCs with GFP and the appearance of fluorescent Nups over time after recombination. A Nup of interest is tagged with a V5 tag followed by loxP sites flanking the selection marker. An eGFP-ORF comes into frame after recombination, so that only protein produced after recombination can be visualized as eGFP-tagged. NUP: nucleoporin open reading frame; V5: V5-tag; loxP: loxP recombination site; eGFP: enhanced green fluorescent protein. Bottom: Cells expressing the indicated Nups endogenously tagged with the RITE cassette were imaged at different timepoints after induction of RITE. Maximum projections are shown. All images are adjusted to the same brightness and contrast values. (B) New and old Mlp1-positive NPCs localize to the nucleolar territory at similar frequency. Percentage of GFP labelled NPCs in the nucleolar territory for Mlp1-RITE(GFP-to-dark) (old) 14 h after induction of RITE and Mlp1-RITE(dark-to-GFP) (new) 45 min after induction of RITE. At least 70 cells were analysed by scoring the localization of NPCs in- or outside the nucleolar territory marked by Nop1-RFP. Blue dots: means of individual biological replicates; error bars: s.d.; ns: not significant by unpaired t-test.

**Figure S5.** Mlp1-truncations localize predominantly to non-nucleolar NPCs. (A) Mean ratio of Mlp1-truncations tagged with GFP in the nucleolar versus non-nucleolar territory. Quantification of data shown in Figure 3B. Blue dots show mean of biological replicates and error bars the standard deviation. At least 50 cells were analysed per replicate and condition. Stars indicate significance in one-way ANOVA with Sidak multiple comparison correction: * p<0.05. (B) Localization of Mlp1ΔNΔC-GFP fragment expressed from a CEN plasmid in the absence of endogenous Mlp1 and Mlp2. Panel shows 2 nuclei with the nucleolus marked by Nop1-RFP. While most cells exhibit low expression and nucleolar exclusion of the fragment (bottom), some cells display high expression and a more homogenous NE staining (top, <5 %). (C) Western blot showing the depletion of Rpb2 by an auxin inducible degron (AID) in the presence or absence of the E3 ligase OsTIR at the indicated timepoints after addition of auxin. Equivalent of 1 OD_600_ of cells loaded per lane. Crm1 was used as a loading control.

**Figure S6.** NPC density in the nucleolar territory increases upon depletion of Nup60. (A) Left: Western blot of the depletion of Nup1 by an auxin inducible degron (AID) in the presence or absence of the E3 ligase OsTIR. Hxk1 was used as a loading control. Equivalent of 1 OD_600_ of cells was loaded in each lane. Right: Spotting showing growth defect of cells expressing Nup1-AID and OsTIR on plates containing auxin. Cells were serial diluted 1:10 and spotted on YPD plates +/-auxin. (B) Western blot of the depletion of Nup60 by an auxin inducible degron (AID) in the presence or absence of the E3 ligase OsTIR. V5 antibody detects both the Nup60-AID as well as the RITE-tagged Nup133. Spontaneous recombination of the RITE tag is visible by the presence of a lower molecular size of Nup133-V5 signal as cells were not grown under selection for the RITE cassette. Crm1 was used as a loading control. Equivalent of 1 OD_600_ of cells was loaded in each lane. (C) Mean ratio of Nup133-RITE(GFP-to-dark) induction in the nucleolar versus non-nucleolar territory upon depletion of Nup60 via the auxin inducible degron. RITE was induced for four hours. At least 50 cells were analysed per replicate and condition. Star indicates significance in one-way ANOVA with Sidak multiple comparison correction: **** p<0.0001. (D) Localization of Nup2-GFP in cells expressing either wildtype Nup60 or the C-terminally truncated version Nup60ΔC. Shown is a single z-slice. Nup2 does not enrich at the nuclear envelope in the absence of its binding site on the C-terminus of Nup60. Exclusion of the soluble protein from the nucleolar region is visible. Nup2-GFP was expressed from a CEN plasmid in the presence of the wildtype copy of *NUP2*. (E) Quantification of Mlp1-GFP foci as shown in Figure 4E. Cells and foci were counted by thresholding on the Nop1-RFP and Mlp1-GFP channels, respectively. At least 200 cells were analyzed per replicate and condition. Stars indicate significance in one-way ANOVA with Sidak multiple comparison correction: **** p<0.0001, ns not significant. (F) Western blot and quantification of Mlp1-GFP expression levels in the indicated mutants. Crm1 was used as a loading control. Equivalent of 1 OD600 of cells was loaded in each lane. No significant (ns) difference was detected by one-way ANOVA with Sidak multiple comparison correction test. All bar graphs: Blue dots show mean of biological replicates and error bars the standard error of mean.

**Figure S7:** Single particle tracking of NPCs in nuclear basket mutants. (A) A 3-state model was fit to tracks obtained from Nup133-RITE(GFP-to-dark) labelled NPCs 14 hours after induction. Bar graphs show the obtained localization errors and diffusion coefficients estimated for the different mutants. The localization errors and diffusion coefficients did not differ significantly between the different strains. Results from individual biological replicates are shown as blue dots, the error bars represent the standard deviation of three biological replicates. At least 1100 tracks were analysed per condition and replicate. P value is from pairwise t-test. (B) A 3-state model with fixed diffusion coefficients (*D*_1_ = 0.001 µm^2^/s, *D*_2_ = 0.005 µm^2^/s, *D*_3_ = 0.03 µm^2^/s) was fit to the same data as in (A). Transition rates between the different states are shown. Error bars represent the standard deviation. P-values are from pairwise t-tests. All other pair-wise comparisons were not significant. Transition rates shown here were used to compute fractions in different states shown in Figure 5D. (C) A 3-state model with fixed diffusion coefficients (*D*_1_ = 0.001 µm^2^/s, *D*_2_ = 0.005 µm^2^/s, *D*_3_ = 0.03 µm^2^/s) was fit to tracks obtained from Nup133-RITE(GFP-to-dark) labelled NPCs expressing Nup60 tagged with an auxin-inducible degron 14 hours after RITE induction. Nup60 degradation was induced with auxin for 90 min. Degradation occurred in strains carrying the OsTIR-E3 ligase. Bar graphs display the fraction of particles in each diffusive state. Results from individual biological replicates are shown as blue dots, the error bars represent the standard deviation. By pairwise t-test, results did not differ between conditions with or without Nup60. At least 1500 tracks were analysed per condition and replicate. (D) Illustration demonstrating the projection of 3D tracks onto a 2D plane. The axes are in µm and the sphere diameter is 2 µm. (E) Quantification of the measured diffusion coefficients from 3D and from projected 2D trajectories. We simulated 200,000 tracks with 11 timesteps moving on a sphere with a diffusion coefficient of 0.01 µm^2^/s. The apparent diffusion coefficient of each track was computed by fitting a first-order polynomial to its mean squared displacements (MSD). Then, tracks were binned according to their mean z position and the average apparent diffusion coefficients were computed for each bin.

**Movie 1.** Tracking of individual yeast NPCs. Nup133-RITE(GFP-to-dark) labelled cells 14 hours after induction of doRITE. Time interval 100 ms. Magenta circles indicate particle detections by Trackmate, red lines show tracks.

## Abbreviations

AID: auxin-inducible degron
doRITE: dilution of RITE-labelled complexes
GFP: green fluorescent protein
NE: nuclear envelope
NPC: nuclear pore complex
Nup: nucleoporin
RFP: red fluorescent protein
RITE: recombination-induced tag exchange
SEM: standard error of the mean
s.d.: standard deviation

## Notes

### Competing Interest Statement

The authors have declared no competing interest.

### Summary of Updates

Revisions in response to reviewer comments: Text changes were made throughout. Added data: 1) Figure S1A: Analysis of RITE switching kinetics by qPCR on genomic DNA shows switching within 1 h. 2) Figure 4A&B: shows contribution of Nup1 to exclusion of NPCs from the nucleolar territory.

