## Supplemental Figures for "Nuclear basket proteins regulate the distribution and mobility of nuclear pore complexes in budding yeast"

### Supplemental Figure S1 - Related to Figure 1

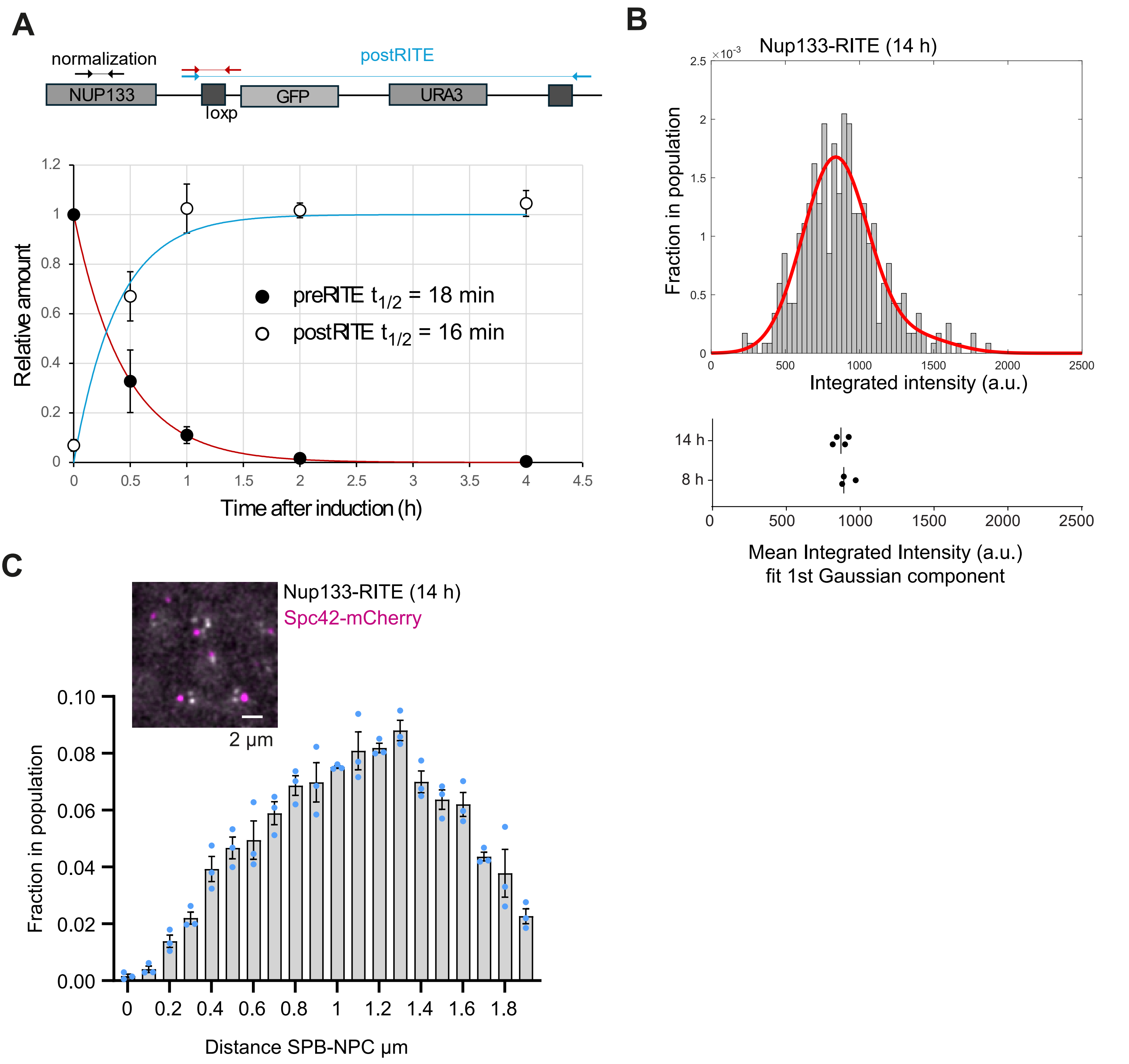


**Figure S1:** doRITE foci show the characteristics of individual NPCs

(A) Kinetics of genomic recombination upon induction of RITE. Quantitative real-time PCR was performed on genomic DNA extracted from Nup133-RITE(GFP-to-dark) cell populations harvested at different time points after RITE induction. Primers detected sequences present before (preRITE, filled circles) or after (postRITE, open circles) recombination. Amounts are shown relative to the NUP133 ORF, which is not affected by the RITE switch. preRITE amounts were normalized to timepoint before induction, postRITE amounts to the 24 hour timepoint. Datapoints represent means from three biological replicates, error bars represent the standard deviation. Red and blue lines are fits with a single exponential, half-times calculated from the fit are given in the legend.

(B) Integrated and background subtracted intensity from individual NPC foci 8 or 14 h after RITE induction. 3D stacks were acquired and intensity was quantified on the best-in-focus slice. Top: Intensity distribution of 350 isolated foci. The red line represents a 2-component Gaussian fit. Bottom: Dots represent the centre peak position of the smaller component of the Gaussian fit for replicates of NPC intensity distributions 8 or 14 hours after RITE induction. Lines represent the median. Each distribution contains >200 individual foci.

(C) NPC-foci do not cluster with SPBs. Distribution of distances of Nup133-RITE(GFP-to-dark) foci at 14 h after RITE induction from the spindle pole body (SPB) labelled by Spc42-mCherry. At least 500 SPB-NPC distances were measured per replicate. Blue dots represent fraction in individual biological replicates. Error bars represent standard error of the mean. Image shows a representative maximum projection of the analysed cells.

### Supplemental Figure S2 - Related to Figure 2

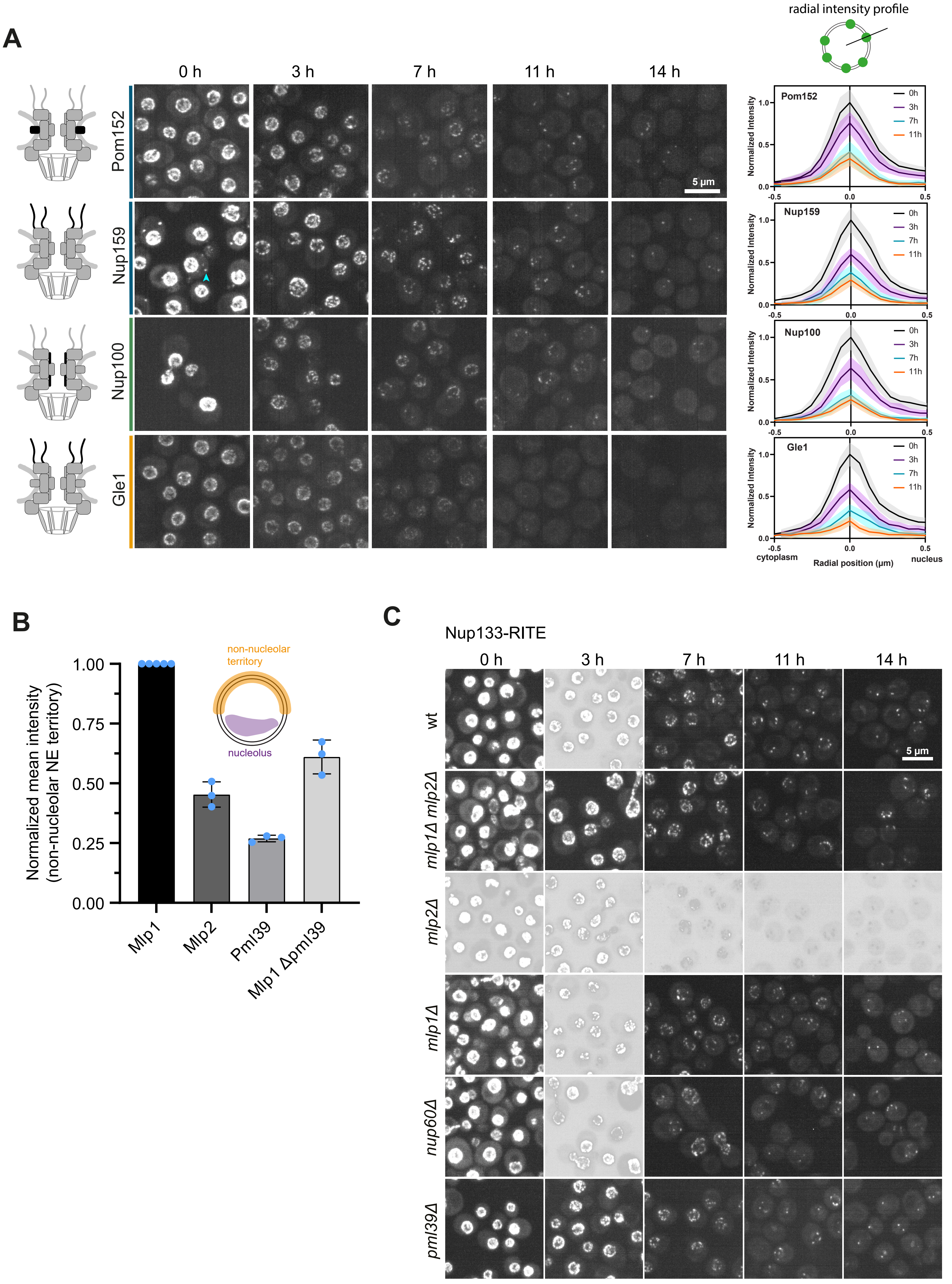

**Figure S2:** Nup dynamics determined by doRITE
(A) Dilution of RITE(GFP-to-dark)-labelled Nups through multiple cell divisions. Cells expressing the indicated Nups endogenously tagged with the RITE cassette were imaged at different timepoints after induction of RITE. Maximum projections are shown. All images are adjusted to the same brightness and contrast values. Cyan arrowhead highlights cytoplasmic Nup159 granule visible only before induction of RITE. Colour bars represent membership to dynamic group in Figure 2A. Schematics on left show location of Nup within the NPC. Graphs on right show radial intensity profile through fluorescent foci on the NE (compare schematic drawing on top) after indicated times of doRITE (solid lines show mean of 20 cells, normalized for 0 h-timepoint, shaded area represents the standard deviation).
(B) The mean intensity of GFP-tagged nuclear basket proteins in the non-nucleolar NE region. Signal outside the nucleus was considered as background, but for the condition Mlp1 Δpml39 the mean signal between nucleoplasm and cytoplasm was considered as background due to higher nucleoplasmic Mlp1 signal. Background subtracted signals are normalized to Mlp1. Blue dots show mean of biological replicates of at least 41 nuclei analysed per condition and replicate. Schematic shows measured area (yellow) of the NE.
(C) Mutations in basket Nups do not affect stability of the NPC scaffold. Cells expressing Nup133 endogenously tagged with the RITE cassette were imaged at different timepoints after induction of RITE. Maximum projections are shown. All images are adjusted to the same brightness and contrast values.

### Supplemental Figure S3 - Related to Figure 2C

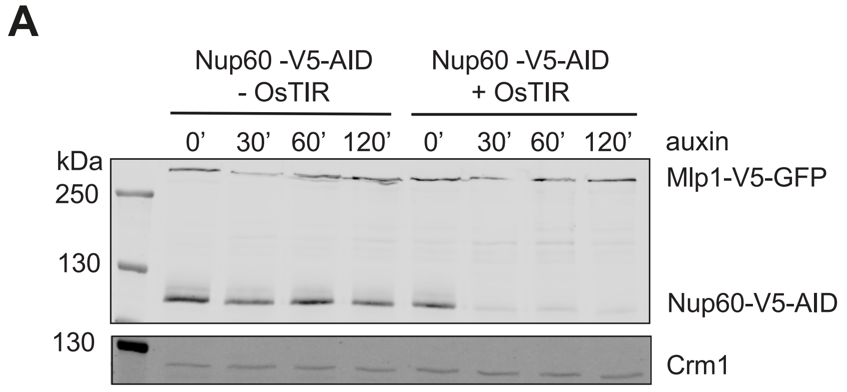


**Figure S3:** Depletion of Nup60 via auxin inducible degradation
(A) Western blot of the depletion of Nup60, in the absence or presence of the E3 ligase adaptor OsTIR, by an auxin inducible degron (AID) at the indicated timepoints after addition of auxin (equivalent of 1 OD600 of cells was loaded in each lane). V5 antibody detects the Nup60-V5-AID as well as the RITE-tagged Mlp1. Crm1 was used as a loading control.

### Supplemental Figure S4 - Related to Figure 3

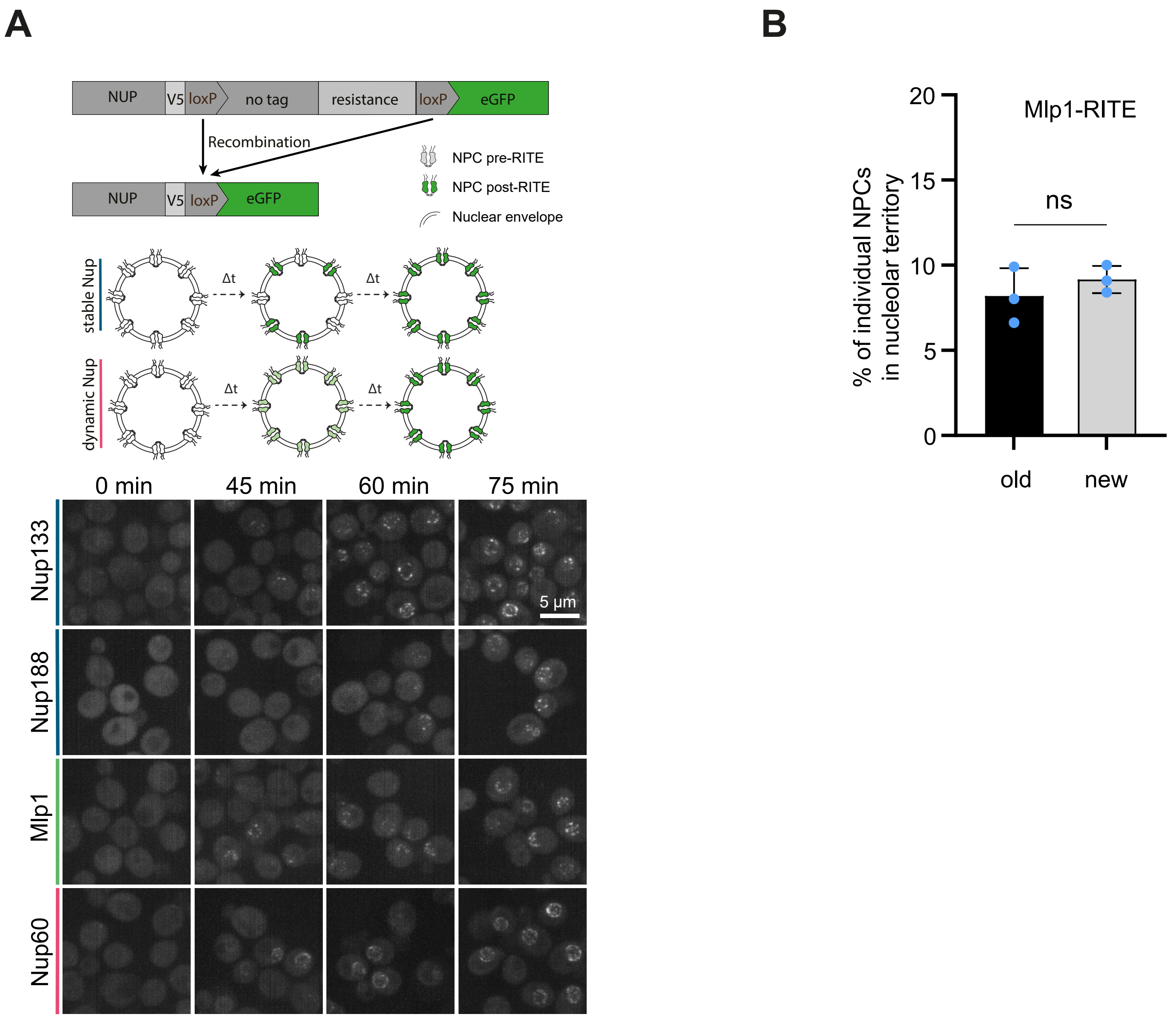


**Figure S4:** A RITE(dark-to-GFP) cassette to visualize newly assembled NPC

(A) Top: Schematic of the dark-to-GFP RITE cassette labelling newly forming NPCs with GFP and the appearance of fluorescent Nups over time after recombination. A Nup of interest is tagged with a V5 tag followed by loxP sites flanking the selection marker. An eGFP-ORF comes into frame after recombination, so that only protein produced after recombination can be visualized as eGFP-tagged. NUP: nucleoporin open reading frame; V5: V5-tag; loxP: loxP recombination site; eGFP: enhanced green fluorescent protein. Bottom: Cells expressing the indicated Nups endogenously tagged with the RITE cassette were imaged at different timepoints after induction of RITE. Maximum projections are shown. All images are adjusted to the same brightness and contrast values.

(B) New and old Mlp1-positive NPCs localize to the nucleolar territory at similar frequency. Percentage of GFP labelled NPCs in the nucleolar territory for Mlp1-RITE(GFP-to-dark) (old) 14 h after induction of RITE and Mlp1-RITE(dark-to-GFP) (new) 45 min after induction of RITE. At least 70 cells were analysed by scoring the localization of NPCs in- or outside the nucleolar territory marked by Nop1-RFP. Blue dots: means of individual biological replicates; error bars: s.d.; ns: not significant by unpaired t-test.

### Supplemental Figure S5 - Related to Figure 3

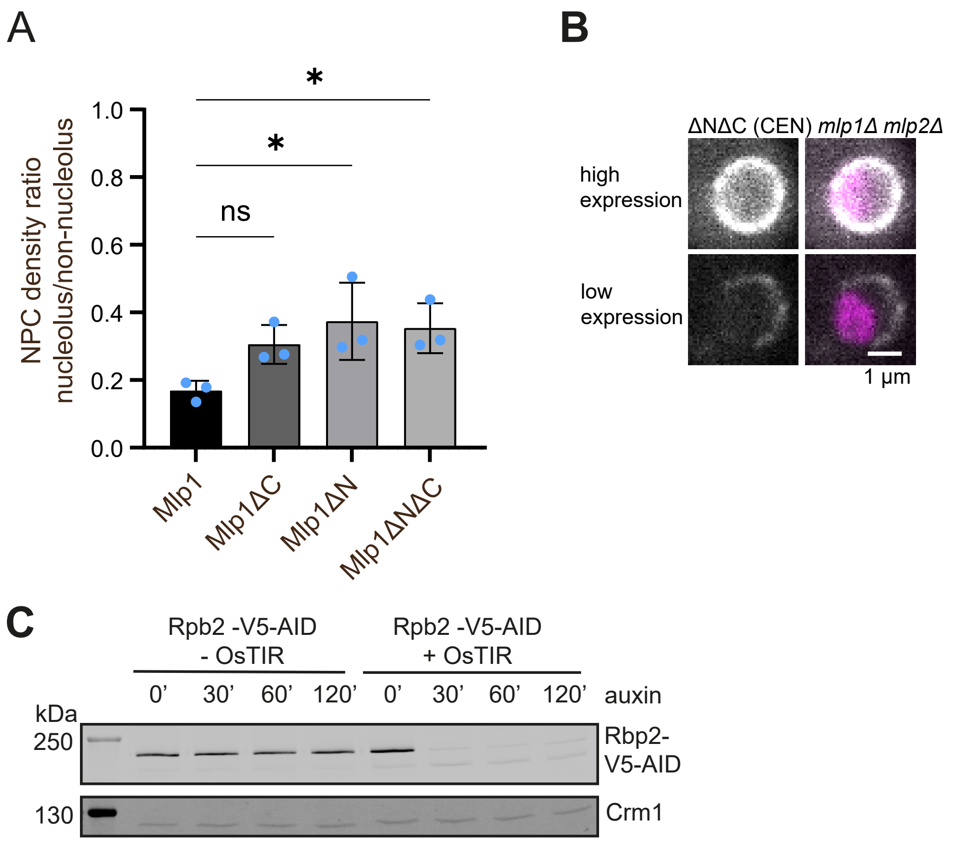


**Figure S5:** Mlp1-truncations localize predominantly to non-nucleolar NPCs
(A) Mean ratio of Mlp1-truncations tagged with GFP in the nucleolar versus non-nucleolar territory. Quantification of data shown in Figure 3B. Blue dots show mean of biological replicates and error bars the standard deviation. At least 50 cells were analysed per replicate and condition. Stars indicate significance in one-way ANOVA with Sidak multiple comparison correction: * $p<0.05$.
(B) Localization of Mlp1ΔNΔC-GFP fragment expressed from a CEN plasmid in the absence of endogenous Mlp1 and Mlp2. Panel shows 2 nuclei with the nucleolus marked by Nop1-RFP. While most cells exhibit low expression and nucleolar exclusion of the fragment (bottom), some cells display high expression and a more homogenous NE staining (top, <5 %).
(C) Western blot showing the depletion of Rpb2 by an auxin inducible degron (AID) in the presence or absence of the E3 ligase OsTIR at the indicated timepoints after addition of auxin. Equivalent of 1 OD600 of cells loaded per lane. Crm1 was used as a loading control.

### Supplemental Figure S6 - Related to Figure 4

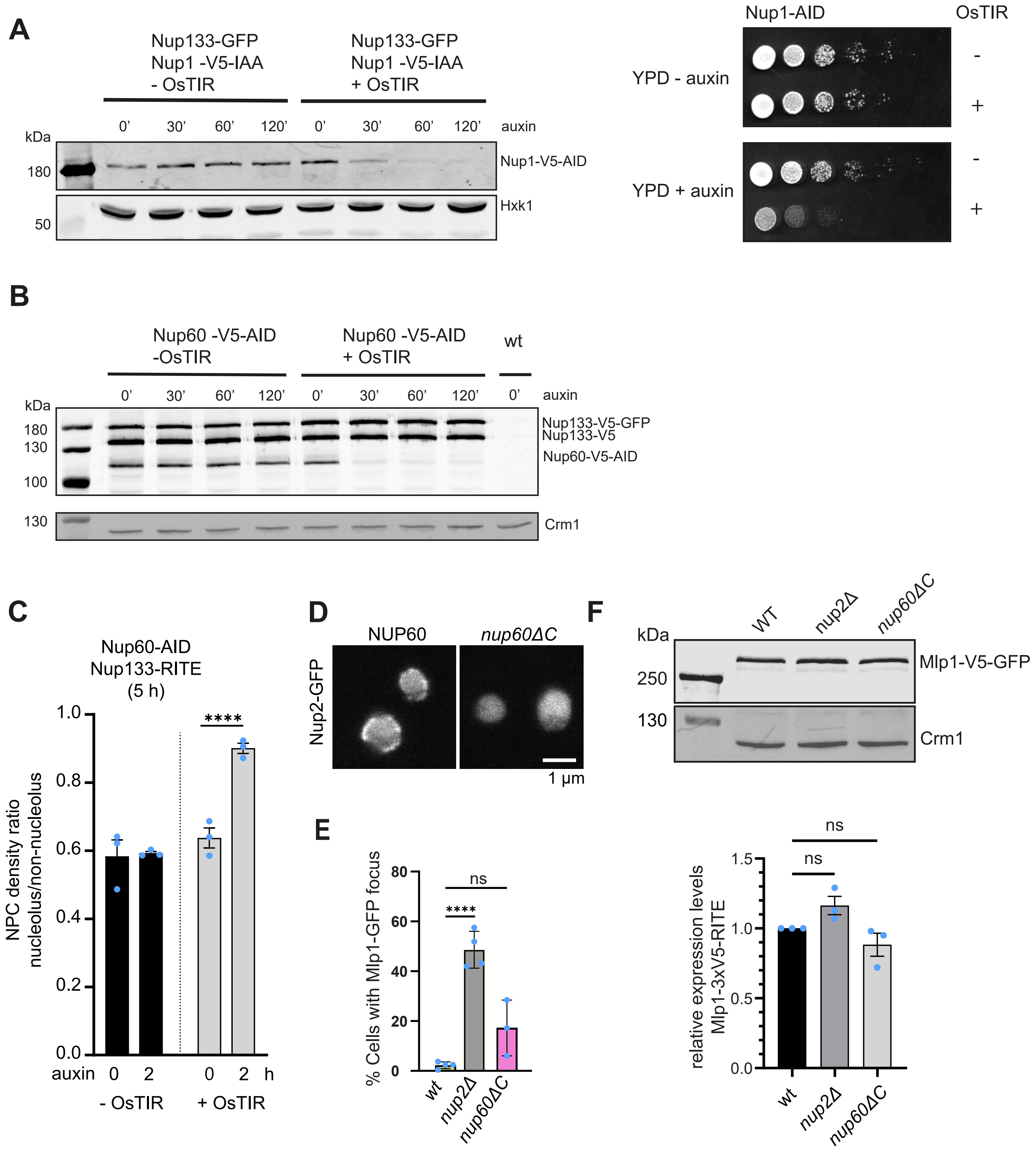

**Figure S6:** NPC density in the nucleolar territory increases upon depletion of Nup60
(A) Left: Western blot of the depletion of Nup1 by an auxin inducible degron (AID) in the presence or absence of the E3 ligase OsTIR. Hxk1 was used as a loading control. Equivalent of 1 OD600 of cells was loaded in each lane. Right: Spotting showing growth defect of cells expressing Nup1-AID and OsTIR on plates containing auxin. Cells were serial diluted 1:10 and spotted on YPD plates +/- auxin.
(B) Western blot of the depletion of Nup60 by an auxin inducible degron (AID) in the presence or absence of the E3 ligase OsTIR. V5 antibody detects both the Nup60-AID as well as the RITE-tagged Nup133. Spontaneous recombination of the RITE tag is visible by the presence of a lower molecular size of Nup133-V5 signal as cells were not grown under selection for the RITE cassette. Crm1 was used as a loading control. Equivalent of 1 OD600 of cells was loaded in each lane.
(C) Mean ratio of Nup133-RITE(GFP-to-dark) induction in the nucleolar versus non-nucleolar territory upon depletion of Nup60 via the auxin inducible degron. RITE was induced for four hours. At least 50 cells were analysed per replicate and condition. Star indicates significance in one-way ANOVA with Sidak multiple comparison correction: **** $p<0.0001$.
(D) Localization of Nup2-GFP in cells expressing either wildtype Nup60 or the C-terminally truncated version Nup60ΔC. Shown is a single z-slice. Nup2 does not enrich at the nuclear envelope in the absence of its binding site on the C-terminus of Nup60. Exclusion of the soluble protein from the nucleolar region is visible. Nup2-GFP was expressed from a CEN plasmid in the presence of the wildtype copy of NUP2.
(E) Quantification of Mlp1-GFP foci as shown in Figure 4E. Cells and foci were counted by thresholding on the Nop1-RFP and Mlp1-GFP channels, respectively. At least 200 cells were analyzed per replicate and condition. Stars indicate significance in one-way ANOVA with Sidak multiple comparison correction: **** $p<0.0001$, ns not significant.
(F) Western blot and quantification of Mlp1-GFP expression levels in the indicated mutants. Crm1 was used as a loading control. Equivalent of 1 OD600 of cells was loaded in each lane. No significant (ns) difference was detected by one-way ANOVA with Sidak multiple comparison correction test.
All bar graphs: Blue dots show mean of biological replicates and error bars the standard error of mean.

### Supplemental Figure S7 - Related to Figure 5

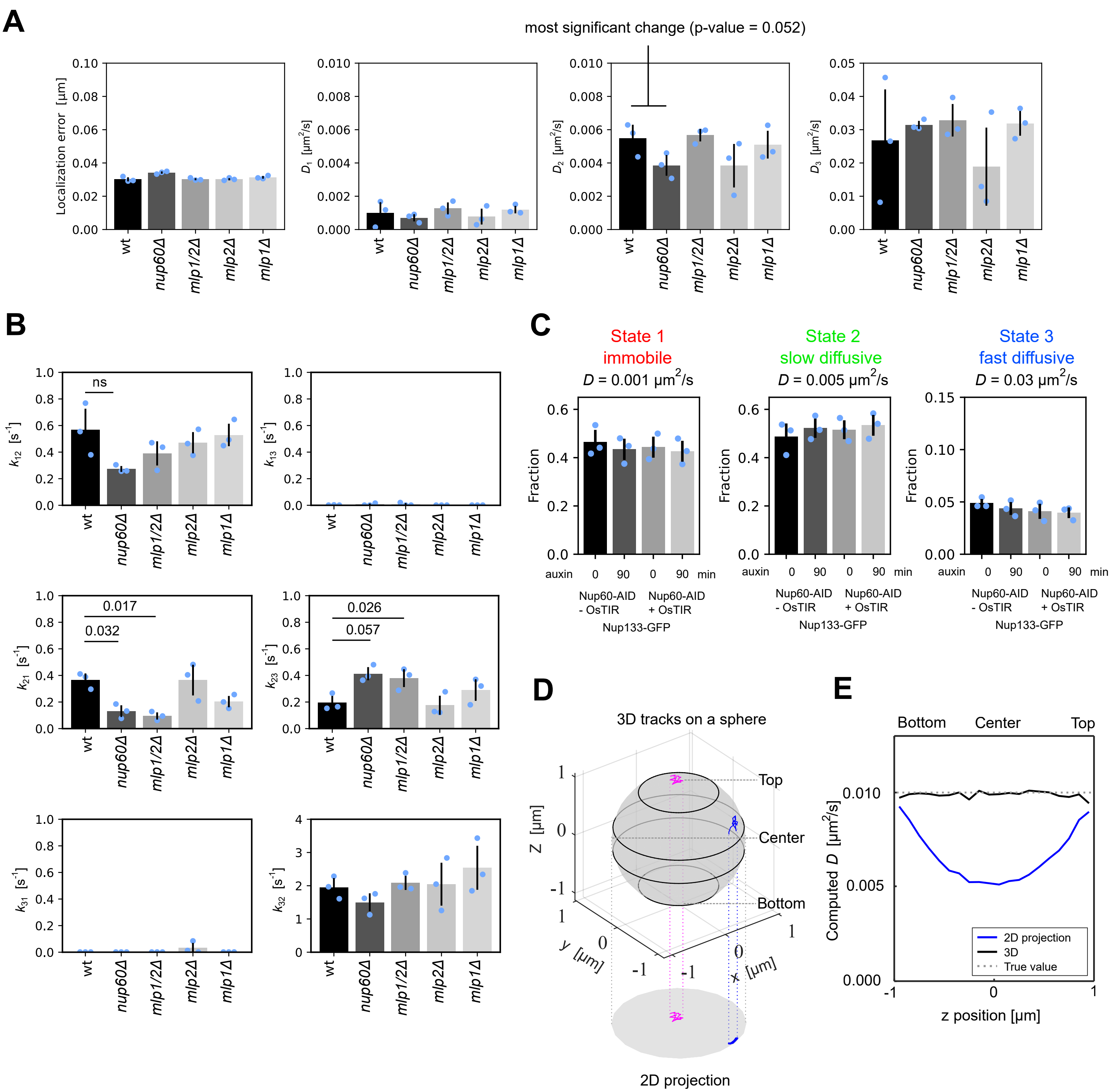

**Figure S7:** Single particle tracking of NPCs in nuclear basket mutants.
(A) A 3-state model was fit to tracks obtained from Nup133-RITE(GFP-to-dark) labelled NPCs 14 hours after induction. Bar graphs show the obtained localization errors and diffusion coefficients estimated for the different mutants. The localization errors and diffusion coefficients did not differ significantly between the different strains. Results from individual biological replicates are shown as blue dots, the error bars represent the standard deviation of three biological replicates. At least 1100 tracks were analysed per condition and replicate. P value is from pairwise t-test.
(B) A 3-state model with fixed diffusion coefficients (D1 = 0.001 µm2/s, D2 = 0.005 µm2/s, D3 = 0.03 µm2/s) was fit to the same data as in (A). Transition rates between the different states are shown. Error bars represent the standard deviation. P-values are from pairwise t-tests. All other pair-wise comparisons were not significant. Transition rates shown here were used to compute fractions in different states shown in Figure 5D.
(C) A 3-state model with fixed diffusion coefficients (D1 = 0.001 µm2/s, D2 = 0.005 µm2/s, D3 = 0.03 µm2/s) was fit to tracks obtained from Nup133-RITE(GFP-to-dark) labelled NPCs expressing Nup60 tagged with an auxin-inducible degron 14 hours after RITE induction. Nup60 degradation was induced with auxin for 90 min. Degradation occurred in strains carrying the OsTIR-E3 ligase. Bar graphs display the fraction of particles in each diffusive state. Results from individual biological replicates are shown as blue dots, the error bars represent the standard deviation. By pairwise t-test, results did not differ between conditions with or without Nup60. At least 1500 tracks were analysed per condition and replicate.
(D) Illustration demonstrating the projection of 3D tracks onto a 2D plane. The axes are in µm and the sphere diameter is 2 µm.
(E) Quantification of the measured diffusion coefficients from 3D and from projected 2D trajectories. We simulated 200,000 tracks with 11 timesteps moving on a sphere with a diffusion coefficient of 0.01 µm2/s. The apparent diffusion coefficient of each track was computed by fitting a first-order polynomial to its mean squared displacements (MSD). Then, tracks were binned according to their mean z position and the average apparent diffusion coefficients were computed for each bin.
